# Enhanced viral infectivity and reduced interferon production are associated with high pathogenicity for influenza viruses

**DOI:** 10.1101/2022.07.29.501947

**Authors:** Ke Li, James M. McCaw, Pengxing Cao

## Abstract

Epidemiological and clinical evidence indicates that humans infected with the 1918 pandemic influenza virus and highly pathogenic avian H5N1 influenza viruses often displayed severe lung pathology. High viral load and extensive infiltration of macrophages are the hallmarks of highly pathogenic (HP) influenza viral infections. However, it remains unclear what biological mechanisms primarily determine the observed difference in the kinetics of viral load and macrophages between HP and low pathogenic (LP) viral infections, and how the mechanistic differences are associated with viral pathogenicity. In this study, we develop a mathematical model of viral dynamics that includes the dynamics of different macrophage populations and interferon. We fit the model to *in vivo* kinetic data of viral load and macrophage level from BALB/c mice infected with an HP or LP strain of H1N1/H5N1 virus using Bayesian inference. Our primary finding is that HP viruses has a higher viral infection rate, a lower interferon production rate and a lower macrophage recruitment rate compared to LP viruses, which are strongly associated with more severe tissue damage (quantified by a higher percentage of epithelial cell loss). We also quantify the relative contribution of macrophages to viral clearance and find that macrophages do not play a dominant role in direct clearance of free virus although their role in mediating immune responses such as interferon production is crucial. Our work provides new insight into the mechanisms that convey the observed difference in viral and macrophage kinetics between HP and LP infections and establishes an improved model fitting framework to enhance the analysis of new data on viral pathogenicity.

**Author Summary:** Infections with highly pathogenic (HP) influenza virus (e.g., the 1918 pandemic virus) often lead to serious morbidity and mortality. HP influenza virus infection is characterised by rapid viral growth rate, high viral load and excessive infiltration of macrophages to the lungs. Despite extensive study, we do not yet fully understand what biological processes leading to the observed viral and macrophage dynamics and therefore viral pathogenicity. Experimental studies have previously suggested that bot viral factors (e.g., viral proteins) and host factors (e.g., the host immune response) play a role to enhance viral pathogenicity. Here, we utilise *in vivo* kinetic data of viral load and macrophages and fit a viral dynamic model the data. Our model allow us to explore the biological mechanisms that contribute to the difference viral and macrophage dynamics between HP and LP infections. This study improves our understanding of the role of interferon on distinguishing immunodynamics between HP and LP infections. Our findings may contribute to the development of next-generation treatment which rely upon an understanding of the host different immunological response to HP influenza viruses.

## Introduction

Influenza is a contagious respiratory disease caused by influenza virus and remains a major public concern [1]. Infections associated with the highly pathogenic (HP) 1918 pandemic H1N1 virus and highly pathogenic avian H5N1 virus often display severe lung pathology, causing fatal infection outcomes in humans [2, 3, 4]. Animal models have been used to understand the mechanisms of viral pathogenicity [5, 6, 7, 8]. High pathogenicity of viruses is often determined by pathogenic outcomes (e.g., the clinical outcomes of infection) in humans [2, 3, 9, 10]. The pathogenicity of influenza virus is not only associated with viral factors (e.g., viral HA protein), but is also influenced by host factors (e.g., the strength of inflammatory response), as reviewed in [11]. For example, although macrophages are important to orchestrate the host immune response, they are also implicated to damage cells through secreted inflammatory cytokines [12, 13, 14, 15]. Some HP viruses can use macrophages as target cells and produce new virus from infected macrophages, altering the antiviral role of macrophages and contributing to viral infection [12, 16, 17]. Perrone et al. compared the outcome of infections with HP and LP strains of two influenza A viruses (i.e., the 1918 pandemic H1N1 virus and an H5N1 virus) in mice and showed that high-pathogenic viruses exhibited a significant higher viral load as early as one day post-infection and a higher number of macrophages in the lungs [18]. However, the temporal dynamics of these viral or host factors, and so the major determinants of the observed differences in viral and macrophage kinetics between HP and LP, remain poorly understood.

Mathematical models have been used to study infection dynamics of influenza virus and its interactions with the host immune response (reviewed in [19]). To explore the potential mechanism(s) leading to the observed difference in viral loads and macrophages between HP and LP infections in the study by Perrone et al. [18], Pawelek et al. fitted a mathematical model to the viral load and macrophage data and found that a higher activation rate of macrophages and an active production of virus by macrophages infected by HP viruses are key drivers leading to higher viral loads and more excessive number of macrophages [20]. More recently, Ackerman et al. [21] fitted a set of mathematical models with different hypothesised mechanisms—leading to distinct immunoregulatory behaviours (e.g., macrophage dynamics)—to strain specific immunological data from [22]. They identified that different interferon production rates are the main causes of variance between infection outcomes in mice infected with low-pathogenic H1N1 or high-pathogenic H5N1 influenza viruses. The two modelling studies provided useful insights into the mechanisms of high pathogenicity and set a framework for assessing other potential mechanisms.

The two modelling studies [20, 21] also left aspects for improvement, both biologically and methodologically. Interferon-mediated immune response, which has been shown to be important for reducing epithelial loss [23], was not considered in [20]. Although the study by Ackerman et al. modelled interferon, they did not compare HP and LP viruses of the same type (rather they compared H5N1 HP vs. H1N1 LP) [21]. Through this study, we aim to identify viral and host factors that determine the observed difference in viral load and macrophage kinetics between HP and LP viruses from same phenotype. Besides, both modelling studies used least-squares method to provide point estimates to model parameters, which may not accurately quantify the uncertainty of estimated parameters and therefore limits our ability to draw reliable conclusions based on parameter estimates [24]. Recent advances in Bayesian statistical inference provides an improved framework for parameter estimation and quantification of uncertainty [25] and can be applied to modelling viral dynamics. We would like to address the above limitations by building an improved framework to study the mechanisms for viral pathogenicity.

In this study, we develop a novel mathematical model which includes macrophage dynamics (i.e., resting, *M*_1_ and *M*_2_ macrophages), interferon-mediated immune response and essential interactions between macrophage and virus. Under a Bayesian statistical framework, we fit the model to available *in vivo* kinetic data for both virus and macrophage populations of both highly pathogenic and paired low pathogenic strains of H1N1 or H5N1 viruses. We use the data-calibrated model to generate and compare a set of metrics that have been used as surrogates for viral pathogenicity [26, 27]. We identify the important role of interferon on distinguishing immunodynamics and the antiviral role of macrophages between HP and LP infections. We also demonstrate that our model reliably captures observed pathogenic behaviours (e.g., the severity of epithelium loss) and provides quantitative estimation of the proportion of damaged cells during HP and LP infections.

## Results

### Severe tissue damage in HP infection

We fit our model to both viral load and macrophage data of HP and LP strains simultaneously under a Bayesian framework (the details of model and statistical implementation, and full diagnostics on the statistical procedures are provided in the Materials and Methods). Model fitting results for H1N1 virus are given in Fig 1. Our model successfully captures the trends of both viral load (Fig 1A) and macrophage number (Fig 1B) for both the HP and LP strains of H1N1 virus and a low level of overlapping of the 95% prediction interval (PI, shaded area) between HP and LP suggests that the quantitative differences in viral load and macrophage are primarily due to different parameter values associated with different strains rather than measurement error. Similar fitting results are observed for infection with the HP and LP strains of H5N1 virus (Figs 1C and 1D).

**Figure 1:**
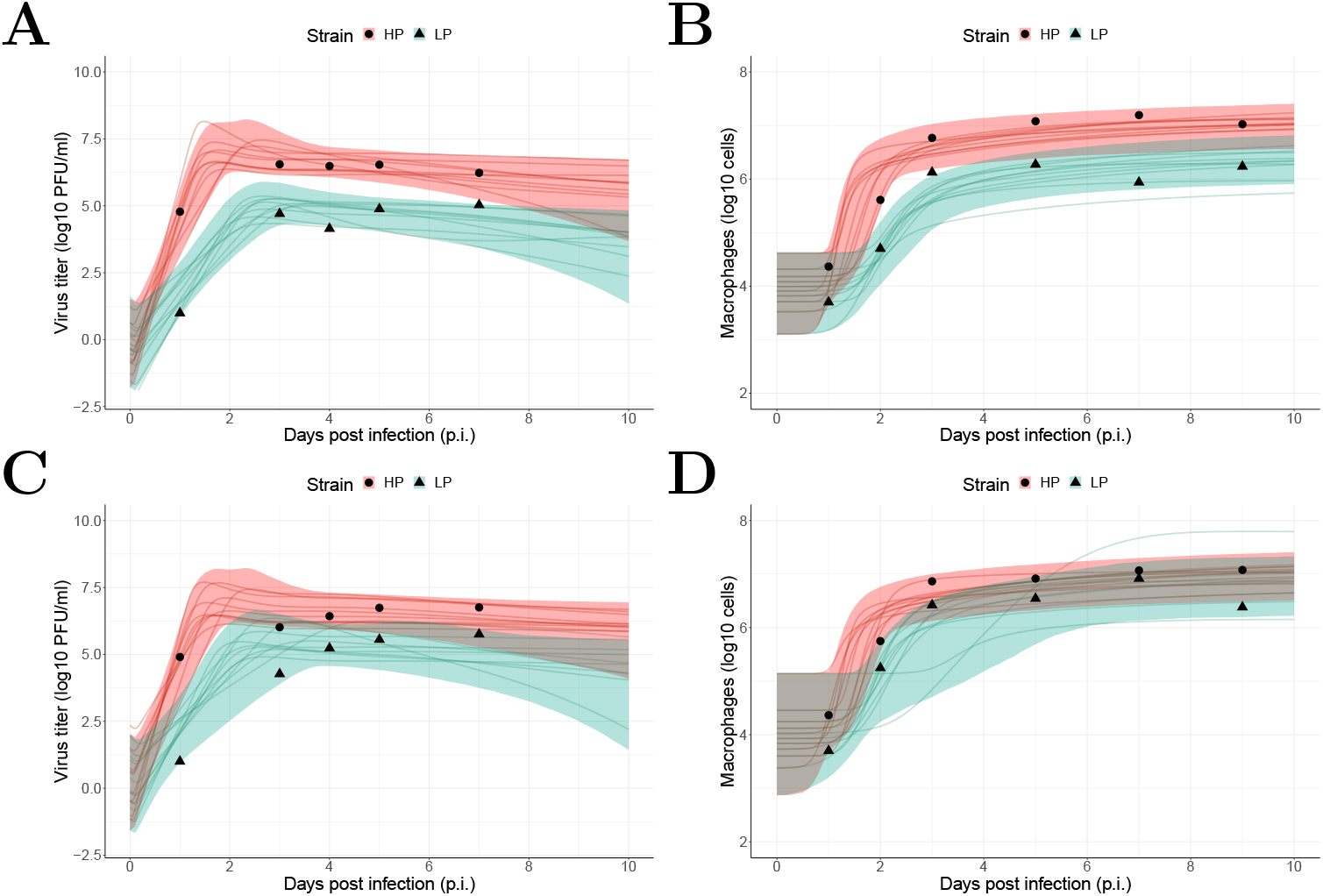
Results of model fitting for virological and macrophage data. Data are presented by solid circles for HP and solid triangles for LP strains. As mentioned in the Materials and Methods, the data were adopted from [18]. We performed 6000 model simulations based on 6000 posterior samples from the posterior distributions of estimated parameters (see SFigs 1 and 2 in *Supporting Information*). (A, B) show a 95% prediction interval (shaded area) of viral load and macrophage for HP (red) and LP (green) strains of the H1N1 viruses, respectively. Solid lines are illustrative viral and macrophage trajectories that are computed based on the basic reproduction number from posterior samples (see Eq. 13 Materials and Methods). (C, D) show the data and model predictions of viral load and macrophage dynamics for HP and LP strains of the H5N1 viruses, respectively.

Using the calibrated model, we then calculate the maximal fraction of epithelium loss (defined by Eq. 14 in Materials and Methods) and the cumulative dead cells (Eq. 15 in Materials and Methods) during infection which are difficult to measure experimentally but are important indicators of infection severity. For H1N1 virus, our model predicts a much larger proportion of epithelial cells (median value 18.4%, 95% predict interval (PI): [3.4%, 97.4%]) are damaged during the HP infection compared to that in the LP infection (median value 0.06%, 95% PI: [0.01%, 0.6%]), as shown in Fig 2A. Similarly for the cumulative number of dead cells shown in Fig 2B, We observe that there is a high cumulation of dead cells (median log_10_(*AUC_D_*) 8.5, 95% PI: [7.7, 8.9]) in the HP infection whereas the cumulation of dead cells is low in LP infection (median log_10_(*AUC_D_*) 6.2, 95% PI: [5.5, 7.1]).

**Figure 2:**
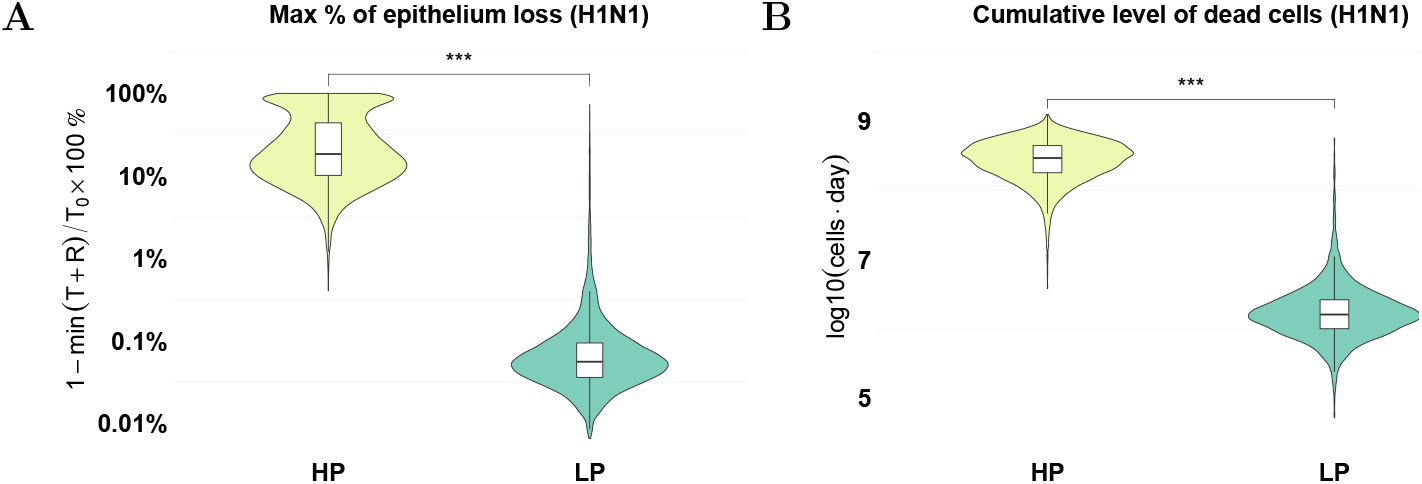
Prediction of tissue damage for H1N1 viruses. The violin plots (coloured) and boxplots (white) give the density and the median and extrema of predicted quantity. (A) model prediction of the maximal epithelium loss for the HP (yellow) and LP (green) strains. (B) model prediction of the cumulative level of dead cells during the infection for both strains. * * *\*p* < 0.001. Calculation formula see Eqs. 14 and 15 in the Materials and Methods. All estimations are computed using 6000 posterior samples from model fitting. The estimations for the H5N1 viruses are given in SFig3 in *Supporting Information*

### A high viral infectivity and a low interferon production contribute to severe tissue damage in HP infection

Given the significant difference in tissue damage between HP and LP virus, we now investigate the underlying biological processes responsible for the differences. We examine the six biological model parameters that may convey the difference between HP and LP virus (i.e., the six parameters assumed to be different between HP and LP in model fitting). To make a direct comparison, we present the ratio of HP’s estimate to LP’s estimate for each parameter in Fig 3 (note that for each parameter there are 6000 ratio values calculated by 6000 paired HP and LP posterior values and thus we show the distribution of the 6000 ratio values in the figure). We observe that for H1N1 the HP strain has a significantly higher viral infectivity *β* (99.7% of the ratio samples are greater than 1 as indicated by dark green. Figs 3B and 3C compare the interferon production rate from infected cells, *q_FI_*, and from activated macrophages, *q_FM_*, respectively. We find that although the HP strain has a decreased *q_FI_*, such that 98.9% ratio samples are lower than 1 (indicated by light green), there is no strong evidence to indicate a difference in *q_FM_*, i.e., approximately half of the posterior estimates for ratios are above 1 (47%) and half below 1 (53%). The results demonstrate that the HP virus is more capable of infecting susceptible cells and reducing interferon response from infected cells. The results are supported by a variety of experimental studies where enhanced infection and replication rates [28, 29] and attenuated interferon production rates [9, 12, 13, 30, 31, 32, 33, 34, 35] are evidenced as possible explanations to high viral pathogenic.

**Figure 3:**
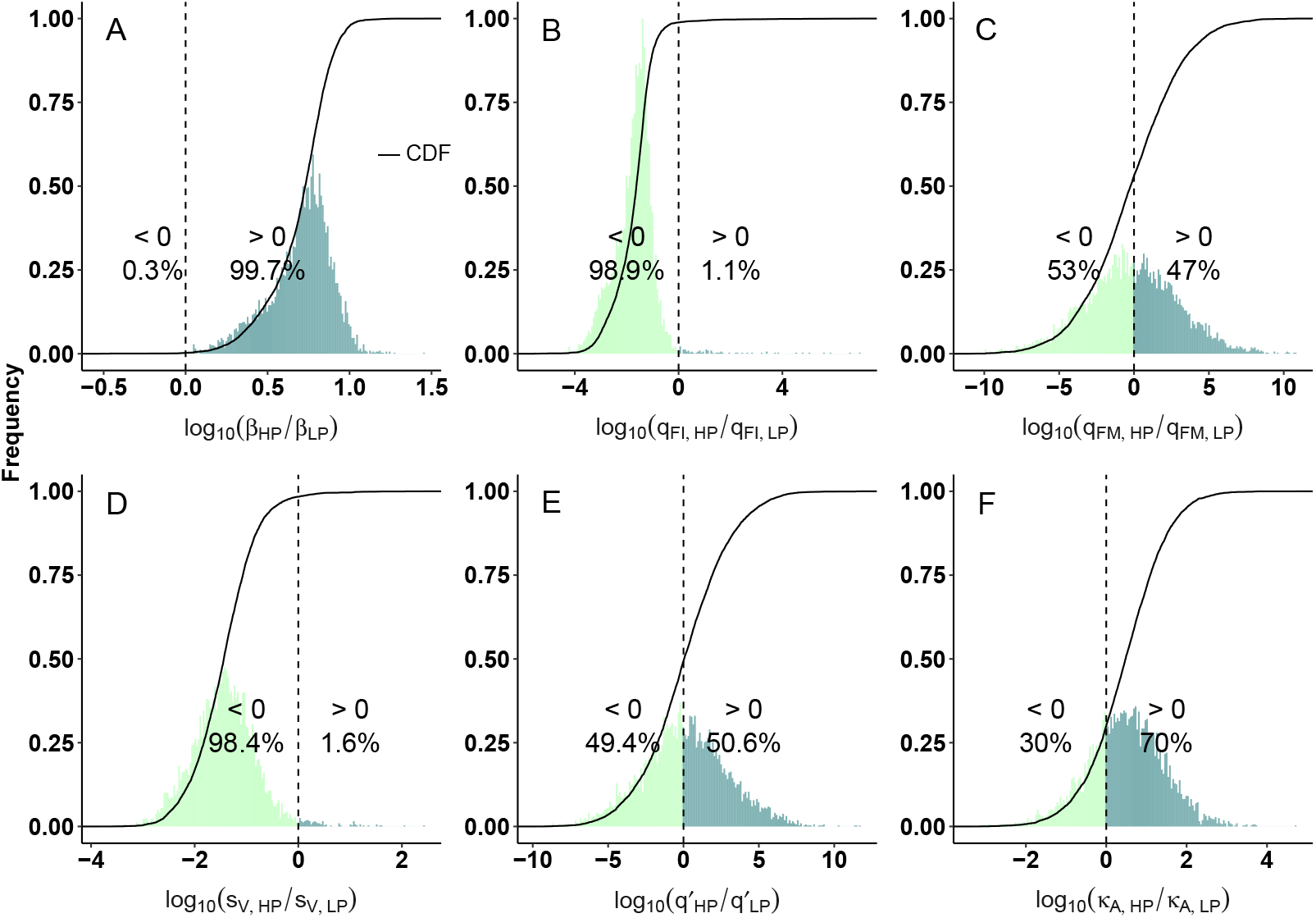
Comparison of estimated model parameters between HP and LP strains of the H1N1 viruses. Histograms show the frequency of the ratios of estimated HP parameters over paired LP model parameters and are normalised to [0,1]. The ratios are presented by distributions of 6000 samples because they are generated by 6000 posterior parameter values. The cumulative density functions (CDFs) are given by the solid lines, and the dashed lines indicate ratios = 0. All ratios are log10-scaled, such that ratios > 0 (dark green) suggest greater values of the HP parameters. Figs (A, B, C) show the ratios of viral infectivity, interferon production rate from infected cells and activated macrophages, respectively. Figs (D, E, F) show the ratios of infection-induced macrophage recruitment rate, macrophage-mediated virus clearance rate and antibody neutralisation rate, respectively. The model parameter comparison for the H5N1 viruses is given in SFig 5 in *Supporting Information*.

Fig 3D shows that the rate of infection-induced macrophage recruitment *s_V_* is lower for the HP strain (98.4% of the ratio samples are less than 1), suggesting that a high recruitment rate is not the cause for the observed high level of macrophages during the HP infection seen in Fig 2. Instead, our model result indicates that high level of macrophages is due to a higher number of infected cells which activate more macrophages. A similar finding was shown by Shoemaker et al. who found that a strong inflammation-associated gene expression occurs once a threshold virus titer is exceeded, demonstrating a strong dependency between the extent of inflammation and the level of virus titer [22].

We further examine how the difference of estimated parameters between HP and LP is associated with the different estimated level of tissue damage shown in Fig 2. We calculate the Partial Rank Correlation Coefficients (PRCCs) between the ratio of estimated parameters and the ratio of epithelium loss between HP and LP strain. We find that the interferon production rate *q_FI_* and infection-induced macrophage recruitment rate *s_V_* are the two leading factors determining the maximum epithelium loss (Fig 4A) and they are negatively correlated with the maximum epithelium loss (PRCC = −0.87 and −0.85 respectively). Analysing the cumulative number of dead cells using the same method, We also find that *q_FI_* and *s_V_* are the two leading parameters driving the difference in the cumulative number of dead cells (Fig 4B), with again negative correlations (PRCC = −0.61 and −0.86). By contrast, the ratio of viral infectivity *β* has a relatively small effect on the ratio of maximum epithelium loss and on the ratio of cumulative number of dead cells. Our results suggest a critical role of interferon in protecting epithelium loss and tissue damage, given *q_FI_* directly determines the rate of interferon production and *s_V_* has an indirect contribution via generating more *M*_1_ macrophages that directly promotes the rate of interferon production (see model equation Eq. 8 in the Materials and Methods). The results are consistent with the earlier finding that interferon can retain a large healthy epithelial cell pool for viral re-infections [23] and supported by Ackerman et al. [21] who found that different interferon production rates are the main causes of variance between infection outcomes in mice infected with low-pathogenic H1N1 or high-pathogenic H5N1 influenza viruses.

**Figure 4:**
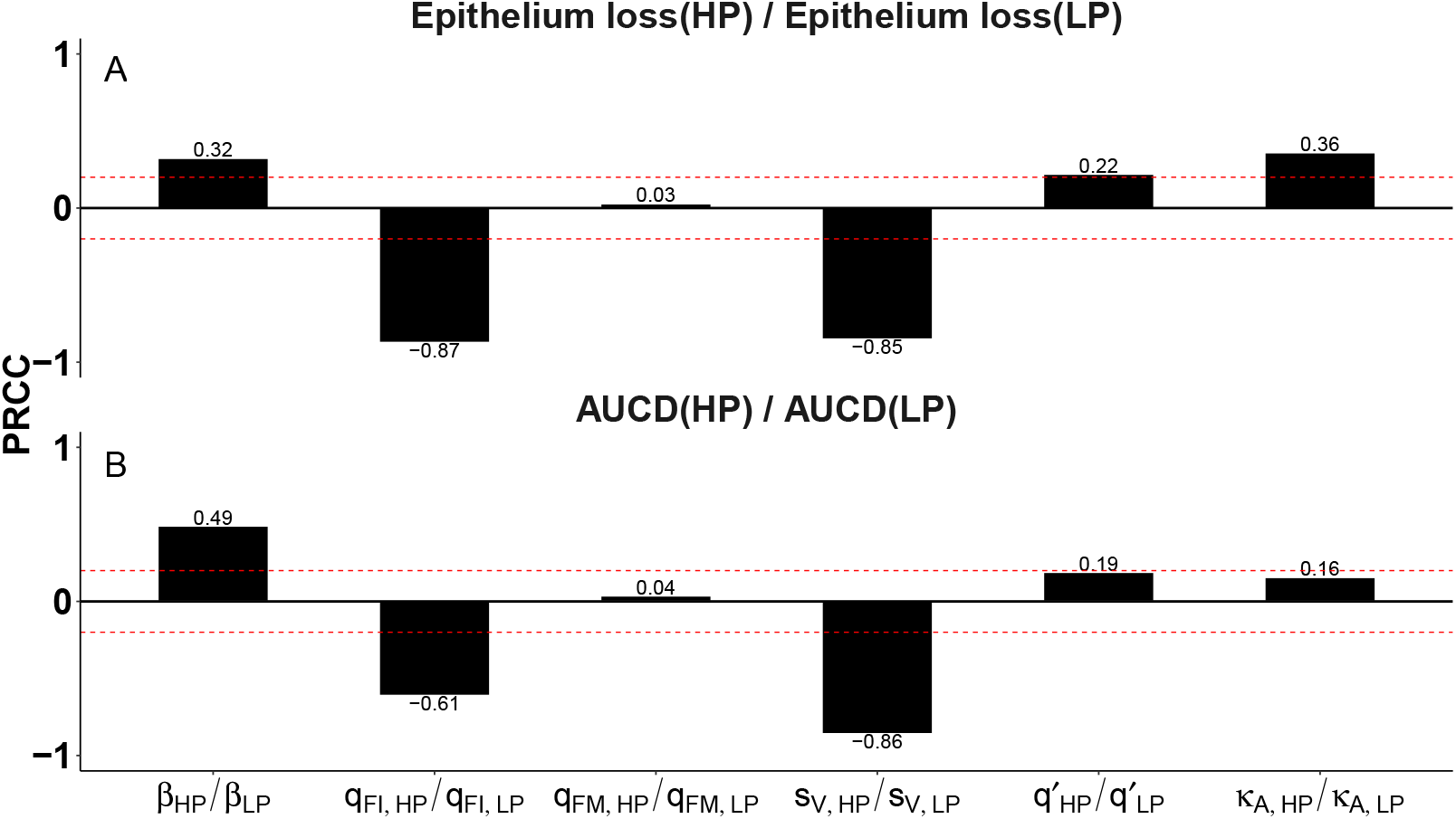
Correlations between estimated model parameters and tissue damage. Partial rank correlation coefficients (PRCC) are calculated with respect to (A) the ratio of max epithelium loss between HP and LP strains, and (B) the ratio of the cumulative dead cells between HP and LP strains of H1N1 viruses. Between the two red dashed lines represents the statistically insignificant values of PRCC. Calculations are based upon 6000 posterior samples from model fitting.

### The role of macrophages on viral clearance

As described in the introduction, the reduced antiviral effect of macrophages may contribute to viral pathogenicity. We here analyse the role of macrophages on viral clearance in both HP and LP infections. In our model, viruses are cleared through three ways: natural decay, macrophage phagocytosis and antibody neutralisation. We use the equation (Eq. 16 in the Materials and Methods) to quantify the contribution of macrophage phagocytosis over the period of infection by a fractional value (e.g., 0 means no contribution and 0.5 means 50% of viral clearance rate is due to macrophage phagocytosis). The prediction interval (PI) can be used to quantify the uncertainty of the contribution fraction. As shown in Fig 5, for H1N1 virus, 95% of model predicted fractions of the contribution of macrophages to viral clearance (indicated by the 90% PI) are below 20% for HP and are below 45% for LP. The upper bounds of the contribution fractions drop significantly for high-confidence range of model predictions, e.g., 60% of of model predicted fractions (indicated by the 20% PI) are less than 0.5% for HP and less than 1.1% for LP. The results indicate the antiviral effects of macrophages is limited in both LP and HP infections, and the relative contribution is even smaller in HP infection. We also compare the relative contribution of macrophages in the HP and LP H5N1 viruses and find a similar result as in the H1N1 viruses (SFig 6 in *Supporting Information*). The result suggests that although macrophages are critical to orchestrate the host immune responses, i.e., initiate and resolve pulmonary inflammation, they are unlikely to be the dominant mechanism to clear free virus.

**Figure 5:**
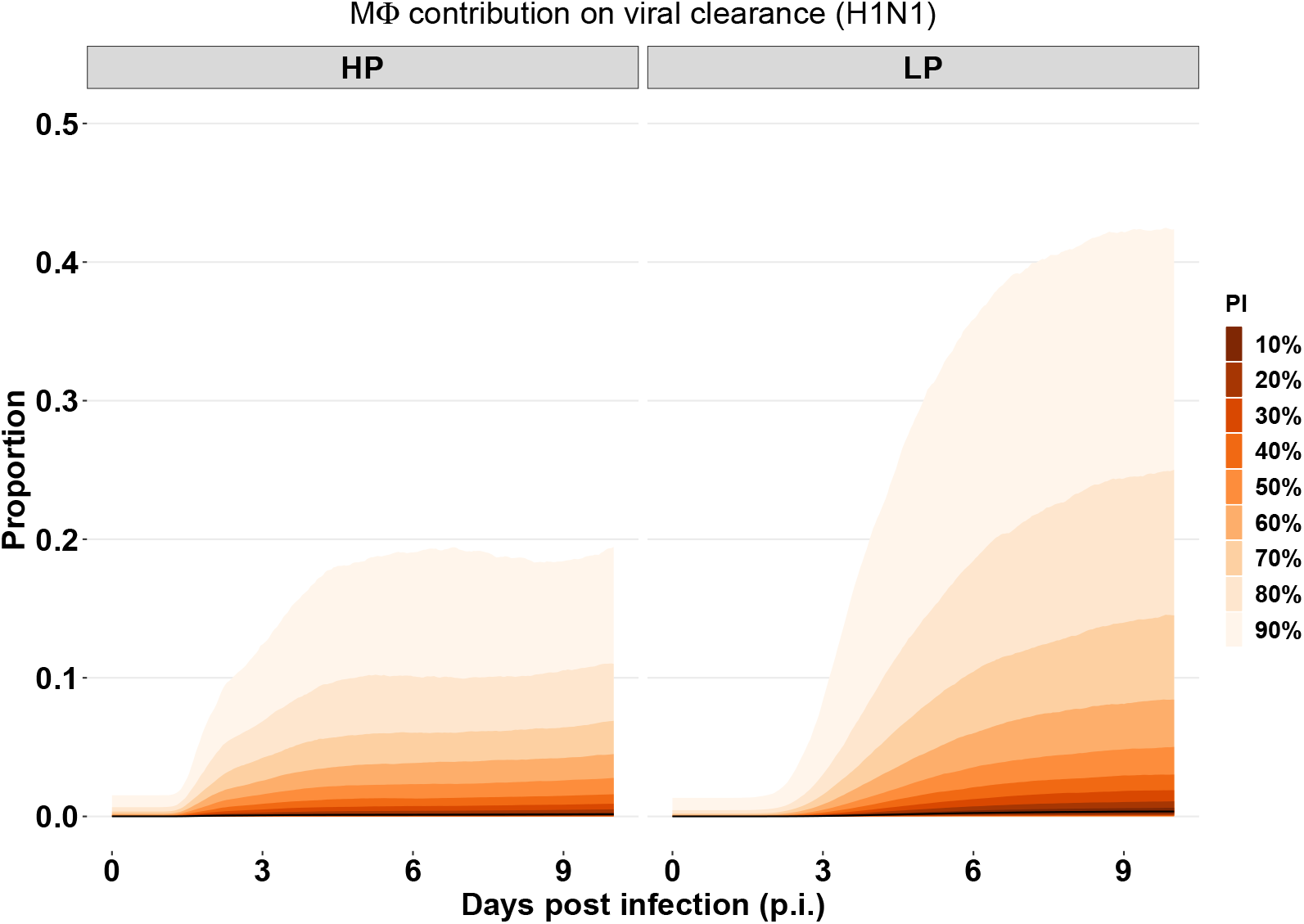
The relative contribution of macrophages on viral clearance in the HP and LP strains of the H1N1 viruses. The prediction interval (PI) is calculated based upon the 6000 posterior samples from model fitting. The median trajectory is indicated by black curve (on the bottom). The predictions for the H5N1 viruses are given in SFig 7 in *Supporting Information*.

### Predicting the effective ways to reduce tissue damage

We have identified three parameters, *β*, *q_FI_* and *s_V_*, that primarily determine the difference in tissue damage (quantified by the maximum fraction of epithelium loss and the cumulative death cell number). This provides insight into the potential targets for the treatment of HP viral infection. Figs 6A, B and C show the impact of varying *β*, *q_FI_* and *s_V_* on the maximal fraction of epithelium loss, respectively. We find that decreasing *β* prevents epithelium loss in both HP and LP infections (Fig 6A). We also observe that increasing interferon production rate *q_FI_* reduces the epithelium loss for the two strains, but the effect is nonlinear (Fig 6B). For example, doubling the production rate halves the epithelium loss, (i.e., epithelium loss is reduced from 30% to 15% for HP and from 0.12% to 0.06% for LP). Reducing 90% of cell loss, however, requires a 10-time increase of *q_FI_*. Furthermore, Fig 6C shows that an enhanced infection-reduced macrophage recruitment rate *s_V_* has almost no influence on epithelium loss for the HP virus. In contrast, it reduces epithelium loss for the LP virus. Note that although the actual magnitude change of epithelium loss is minor for the LP strain, the percentage change is comparable between the two strains. Figs 6D, E and F show the dependency of cumulative death cell number upon *β*, *q_FI_* and *s_V_*, respectively. We find the cumulative death cell number is sensitive to all three parameters for both HP and LP strains. The results imply that the maximal epithelium loss and the cumulative level of dead cells are strongly associated with *β* and *q_FI_*, and reducing viral infectivity or boosting interferon production can prevent epithelium loss. The results also suggest enhancing macrophage recruitment rate *s_V_* during infection can reduce the dead cell accumulation.

**Figure 6:**
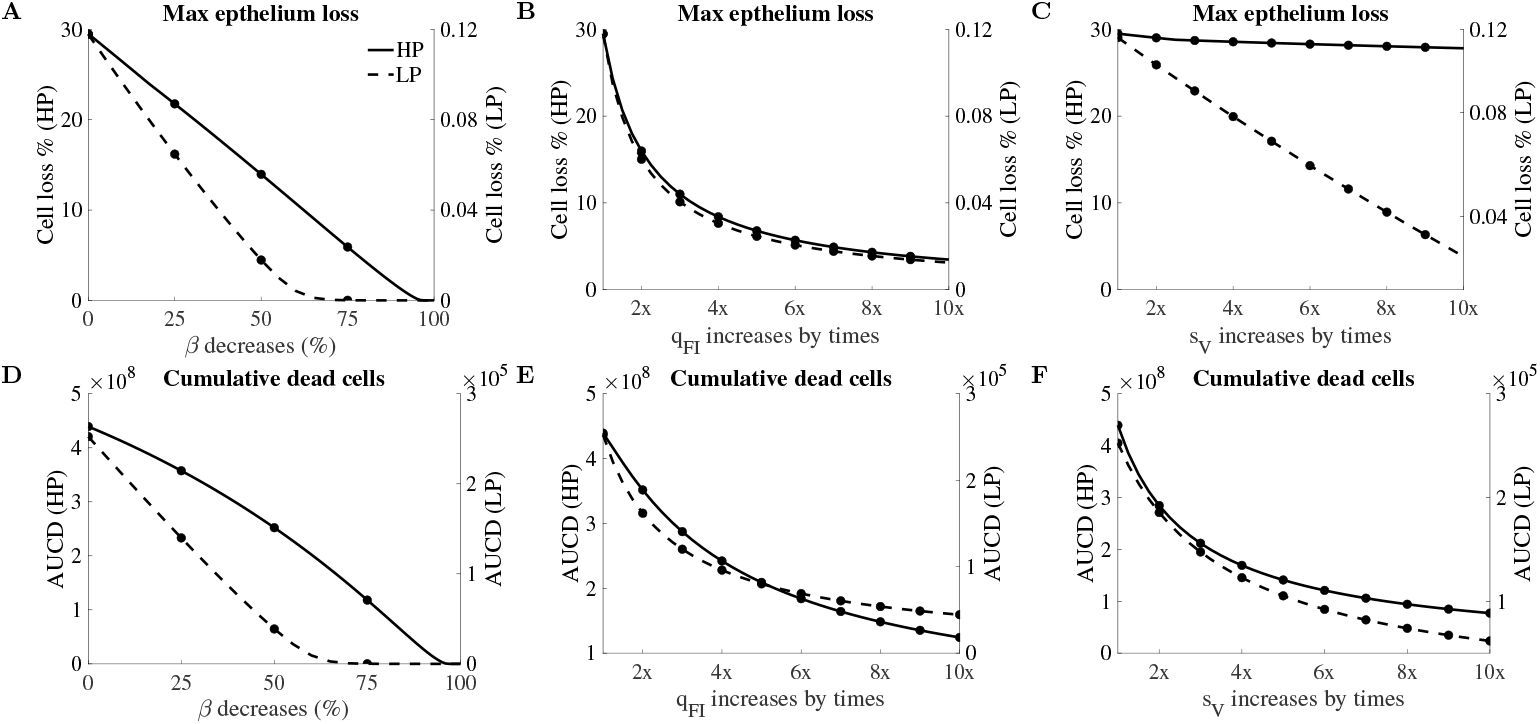
Parameter driving tissue damage for the H1N1 viruses. Solid lines are for the HP and dashed lines are for the LP strains. Figs (A, B, C) give the sensitivity analyses of the impact of *β*, *q_FI_* and *s_V_* on maximal epithelium loss. Figs (D, E, F) show the impact of the same three model parameters on the cumulative dead cells.

**Figure 7:**
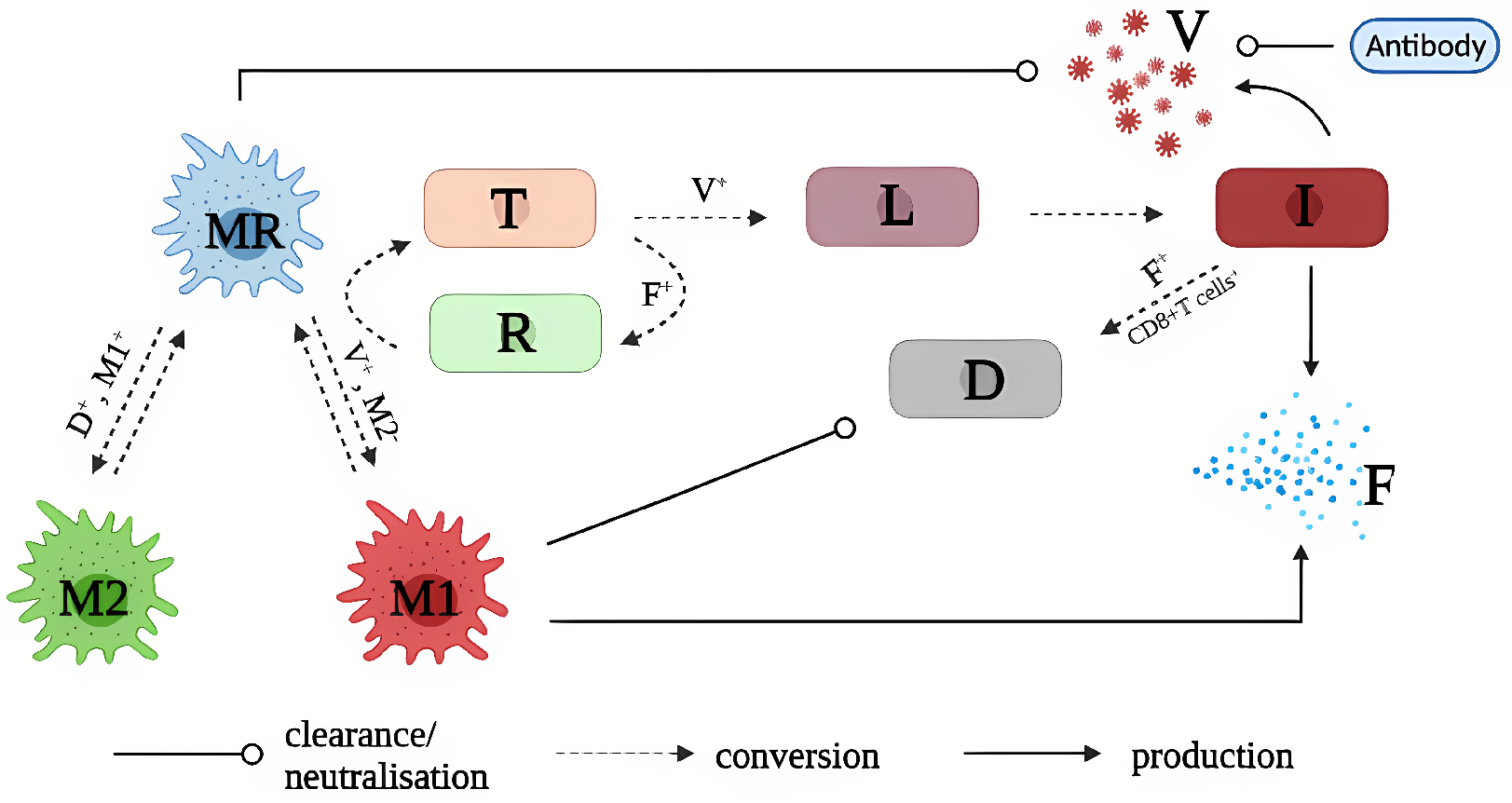
A model diagram of immune response to influenza viral infection. Detailed model (Eqs. 1–10)) description is given in Materials and Methods. Plus (+) superscript indicates the promotion of a biological process, and minus (–) superscript means the inhibition of a process. In brief, influenza virus (*V*) turns susceptible epithelium cells (*T*) into eclipse-phase infected cells (*L*) which in turns, become infected cells (I) that actively produce new virus. Virus also infects resting macrophages (*M_R_*) and turns them into pro-inflammatory macrophages (*M*_1_). Virus is cleared through the *M_R_* macrophage ingestion and antibody neutralisation. Infected cells (*I*) and *M*_1_ macrophages produce interferons (*F*) that turns susceptible cells (*T*) into refractory cells (*R*). The refractory cells (*R*) lose protection and turn back to *T*. Infected cells (*I*) are killed and become dead cells (*D*) through interferons- and CD8^+^ T cells-mediated clearance. *M*_1_ macrophages clear dead, which facilitates the conversion of *M_R_* to anti-inflammatory *M*_2_ macrophages. Both activated *M*_1_ and *M*_2_ macrophages convert back to *M_R_* macrophages at certain rates. For clarity, flows depicting the natural decay of activated macrophages (*M*_1_ and *M*_2_), virus (*V*) and interferons (*F*), and the replenishment of resting macrophages (*M_R_*) and target cells (*T*) are not showed in the diagram.

## Discussion

In this work, we identified biological mechanisms that are associated with high pathogenicity of *in vivo* H1N1 and H5N1 infections through fitting a viral dynamic model to experimental data under a Bayesian framework. Our findings support and contribute to the current knowledge that is relevant to two frequently studied experimental explanations on the drivers of high pathogenicity for influenza viruses (i.e., a higher viral infectivity and a reduced interferon response). Estimated marginal posterior densities of model parameters demonstrate that HP viruses have enhanced viral infection rates (i.e., higher *β*) and reduced interferon production rates (i.e., lower *q_FI_*) compared to LP viruses. Our estimation results also explain the difference in viral and macrophage kinetics between HP and LP infections. As shown by previous studies [23, 36, 37], a higher viral infection rate leads to a faster viral growth and an attenuated interferon production leads to a higher peak viral loads.

Our work quantified the difference of tissue damage between HP and LP infections. We predicted a larger proportion of epithelium loss and a high level of dead cells are caused in HP infections (Fig 2 for H1N1 and SFig 3 for H5N1 in *Supporting Information*). Our model predictions—a high fraction of epithelium loss and a high level of dead cells in HP infection—are supported by clinical evidence. Severe destruction of lung tissue [2] and severe tissue consolidation with unique destruction of the lung architecture [2, 38] have been seen in patients infected with HP influenza viruses, leading to lung pathology [28, 39, 40, 30, 41]. The severity of tissue damage also resulted in different mechanisms of viral resolution. While target cell depletion remains a mechanism to limit viral replication in HP infections, a timely and strong activation of immune response explains viral resolution in LP infections (SFig 4 in *Supporting Information*). As shown by Cao and McCaw, the mechanisms for viral control can strongly influence the predicted outcomes of antiviral treatments [42]. For example, different viral dynamics (e.g., long-last infection or chronic infection) were observed in response to an increasing drug efficacy when target cell depletion is a mechanism for viral resolution. In contrast, a consistent viral behaviour (i.e., an early clearance and a shorter infection) was observed when drug efficacy increased in an immune response driven viral resolution model. Therefore, the analysis of the influence of antiviral treatment on HP and LP infections is a promising future direction based on our work.

Using a Bayesian statistical method, our modelling work demonstrated that high virulence of H1N1 and H5N1 viruses, and our estimation provided evidence to previous experimental work. Although our work identified HP and LP viruses differ in viral infectivity and interferon production rate, we cannot (and do not attempt to) rule out other possible mechanisms or drivers of high pathogenicity proposed in literature. For example, production of virus by infected macrophages could be an important factor influencing viral pathogenicity [17], although there is conflicting evidence on whether macrophages can be productively infected by influenza virus [15, 16, 43]. The abortive or productive infection of macrophages may also be strain-dependent and/or macrophage-dependent (i.e., resident or monocyte-derived macrophages) [17]. Thus, we have not explicitly investigate this mechanism in our study.

Viral dynamical models are particularly useful in the quantification of modelled biological processes by fitting to experimental data [19]. In this work, we fit our model to both viral load and macrophage data to estimate model parameters. Although consistent with earlier studies where both viral load and macrophages were used in model fitting, the effect of incorporating macrophage data into model fitting remains unclear. Using a simulation-estimation method, we showed that macrophage data provides valuable information on parameter estimation, reducing the uncertainty of predicted time series of macrophages and estimates of the recruitment rates of macrophages (i.e., *s_M_* and *s_V_*). By contrast, viral load data alone are insufficient to reliably recover macrophage dynamics (see S3 Text in *Supporting Information*). Macrophages have been shown to clear viruses by internalisation and lysosomal degradation [44, 45], but their relative contribution to viral clearance compared to other pathways has not been quantified. Our model predicted the contribution of macrophages on viral clearance (among all the modelled mechanisms for viral clearance) is relatively small in both HP and LP infections of H1N1 (Fig 5) and H5N1 (SFig 6 in *Supporting Information*) viruses, suggesting that macrophages may not play a dominant role in direct clearance of free virions. Our model also suggests that the relative contribution of macrophage to viral clearance in HP viral infection is smaller than that in LP infection. This is because resident macrophages (*M_R_*) do not replenish during HP infection while they can quickly replenish in LP infection (SFig 7 in *Supporting Information*), which increases the available macrophages to participate in viral clearance. Another mechanism [15] related to productive replication of HP viruses in macrophages has been to have significant consequences for the antiviral functions of macrophages, as reviewed in [17].

Our study has some limitations. Rather than explicitly modelling the dynamics of CD8^+^ T cells and antibodies [36, 46], we used hill functions to capture their dynamics. We assumed the adaptive immune response dominates infected cell or viral clearance at day 5 post infection regardless of macrophage dynamics. Macrophages, however, have been shown to act as antigen presenting cells and mediate the activation of different arms of adaptive immunity. For example, *M*_1_ type macrophages help to activate the cellular adaptive immune response whereas *M*_2_ type macrophages contribute to the activation of humoral adaptive immunity [47, 48]. Extension of the model to include the interactions between different populations of macrophages and adaptive immunity is important but requires additional data on the adaptive immune response for both HP and LP, which are not immediately available in the literature. Another limitation is that we did not estimate conversion rates between different populations of macrophages, such as *k*_1_ and *k*_2_, due to a lack of detailed macrophage kinetic data. As a result, the kinetics for each specific macrophage population could not be calibrated against data. The interactions among macrophage populations, e.g., the rate of conversion from one type to another, could be an important factor to understand influenza disease severity. In future work, our model can be used to estimate the relevant parameters and predict detailed macrophage dynamics given availability of data of different macrophage populations.

## Materials and Methods

### Mathematical Models

In this study, we incorporated a dynamic model of macrophages into a viral dynamic model. The model explicitly considered the conversion among different populations of macrophages, essential interactions between virus and macrophages, and different arms of immune responses. The model is described by a set of ordinary differential equations (ODEs).

Eqs. 1–3 describe the detailed macrophage dynamics. In the absence of viral infection, we assume all macrophages are resting macrophages (*M_R_*), and *M_R_* is assumed to have a constant supplementary rate and decay rate at *s_M_* and *δ_MR_* per day, respectively. Thus, the number of macrophages is stable at homeostasis, such as 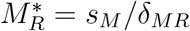 in a disease-free condition. In the presence of viral infection, influenza virus acting as a perturbation to macrophage dynamics, activates *M_R_* macrophages, turning them into pro-inflammatory macrophages *M*_1_ at a maximal rate *k*_1_. The activation is influenced by viral load (*V*/(*V* + *V*_50_)) and regulated by anti-inflammatory *M*_2_ macrophages (1/(1 + *αM*_2_)). Activated *M*_1_ macrophages convert back to the resting macrophages or decay at constant rate *k*_−1_ and *δ_MA_* per day, respectively. The *M*_2_ macrophages regulate the activation of *M*_1_ macrophages to avoid excessive inflammatory response [49]. *M*_1_ macrophages phagocyte apoptotic and dead cells, producing regulatory cytokines (not explicitly modelled), which is represented by *M*_1_*D*/(*D* + *D*_50_). In the presence of these cytokines, resting macrophages *M_R_* convert to *M*_2_ macrophages at a maximal rate *k*_2_. Activated *M*_2_ macrophages decay or convert back to the resting state at constant rates *δ_MA_* and *k*_−2_, respectively.

Eqs. 4–7 describe the interaction between virus and epithelial cells, and between virus and the host immune responses. In detail, epithelial cells (*T*) are infected by influenza virus (*V*) and become latent-state infected cells (*L*) which do not produce new viruses at an infectivity rate *βV* per day. The susceptible epithelial cells are protected and convert to refractory cells (*R*) in the presence of interferon (*F*) at a rate *ϕF* per day, and refractory cells convert back to susceptible cells at a rate *ξ_R_*. We also assume susceptible cells are replenished at a rate *g_T_*(*T* + *R*)(1 − (*T* + *I* + *R*)/*T*_0_), where *T*_0_ is the maximal number of epithelial cells that line the upper respiratory tract. Infected cells in eclipse phase convert to infected cells (*I*) that actively produce virus at a rate ℓ per day. Three mechanisms are considered for the clearance of infected cells (*I*), such as natural decay at a constant rate *δ_I_* per day; interferon-mediated clearance at a rate *κ_F_F* per day, and CD8^+^ T cells mediated infected clearance at a rate 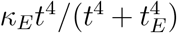 per day. Note that we do not explicitly model the dynamics of CD8^+^ T cells. A hill function is used to represent the activation of adaptive immunity, we set *t_E_* as 5 so that CD8^+^ T cells only play a significant role after day 5 post infection as showed in [50]. New virus is produced by *I* at a rate *p_I_ I* viruses per day. The decrease of virus is either due to natural decay, macrophage-mediated phagocytosis or antibody neutralisation at a rate 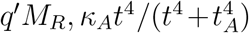 per day, respectively.

Eqs. 8–9 describe the one of the interferon dynamics and the dynamics of refractory cells. We assume Interferon (*F*) is produced either by infected cells (*I*) or macrophages (*M*_1_) at a rate *q_FI_I* or *q_FM_M*_1_ unit of interferons per day, respectively, and decay rate a rate *δ_F_* per day. The dynamics of dead cells (*D*) is described by Eq. 10. Cleared infected cells (*I*) become dead cells (*D*) through *δ_I_I, κ_F_F* and 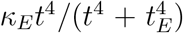, and dead cells is removed from the system either due to natural decay at a rate *δ_D_* per day or killed by macrophages *κ_D_M*_1_ per day.

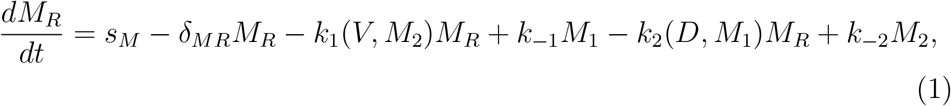

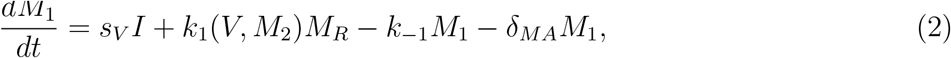

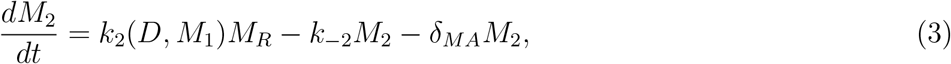

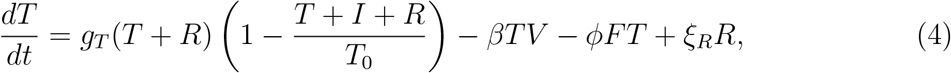

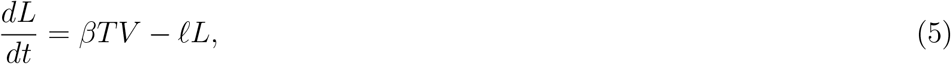

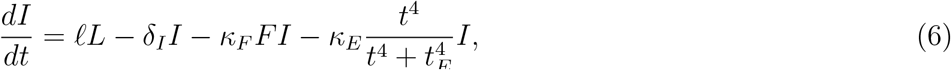

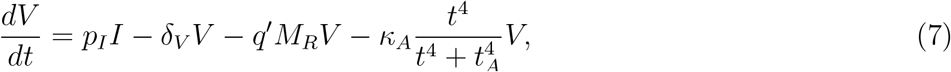

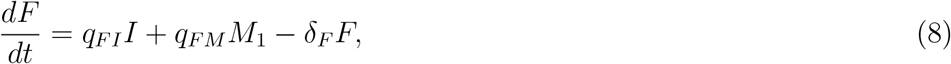

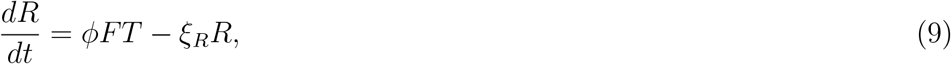

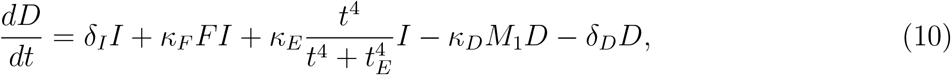

where 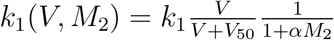 and 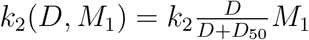.

### Statistical Inference

*In vivo* kinetic data of both virus and macrophage population were extracted using WebPlotDigitizer (version 4.4) from [18]. Female BALB/c mice were intranasally infected with HP (A/1918 H1N1 and A/Thailand/16/2004 H5N1) and LP (A/Texas/36/91 and A/Thailand/SP/83/2004) influenza viruses, and lungs were harvested for viral load and macrophage measurement at various time points post infection. Three mice were measured per time point for infection with each viral strain.

We applied a Bayesian inference method to fit the dynamic model (detailed in Mathematical Models) to the log-transformed virological and macrophage data. In detail, we use the model to estimate 8 parameters, and the parameter space is denoted as 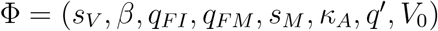. Upon model calibration, we fixed all other parameters to previous estimated values in the literature. We fixed the parameter values because the experimental study [18] does not provide sufficient data for parameter estimation. The fixed parameter values are given in S2 Text in *Supporting Information*.

We assumed HP and LP viruses differ in 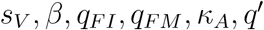 but have same *s_M_* and *V*_0_. This is a reasonable assumption given inbred mice having similar number macrophages in the absence of infection (i.e., same *s_M_*), and inoculation size is same for HP and LP infection (i.e., same *V*_0_). The prior distribution for the estimated model parameters is given in S2 Text in *Supporting Information*. The distribution of the observed log-transformed viral load and macrophage data is assumed to be a normal distribution with a mean value given by the model simulation results and standard deviation (SD) parameter with prior distribution of a normal distribution with a mean of 0 and a SD of 1.

Model fitting was performed in R (version 4.0.2) and Stan (Rstan 2.21.0). Samples were drawn from the joint posterior distribution of the model parameters using Hamiltonian Monte Carlo (HMC) optimized by the No-U-Turn Sampler (NUTS) (see [25] for details). In particular, we used three chains with different starting points and ran 3000 iterations for each chain. The first 1000 iterations were discarded as burn-in, and we retained 6000 samples in total from the 3 chain (2000 for each). Detailed diagnostics and results can be found in S1 Text in *Supporting Information*.

### Model prediction

The model prediction for any quantities *z* and data *y* given parameter set *θ*, we compute

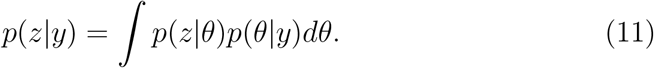

Here, quantities z are viral reproduction number, the maximal epithelium loss and the cumulation of dead cells. The effective reproduction number of viral replication (*R_t_*) is given by

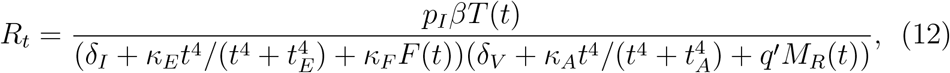

where *T*(*t*), *F*(*t*) and *M_R_*(*t*) are the number of susceptible of epithelial cells, interferons and resting macrophages during infection. The killing effect of CD8^+^ T cells and the neutralization effect of antibodies are represented by 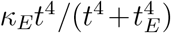 and 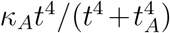, respectively. At *t* = 0, *T*(0) = *T*_0_, *F*(0) = 0 and *M_R_*(0) = *s*/*δ_MR_*, and *R*_0_ is called the basic reproduction number of viral infection, which simplifies to

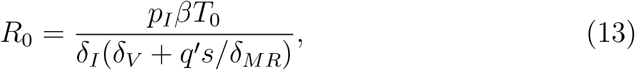

The maximal % of epithelium loss is given by

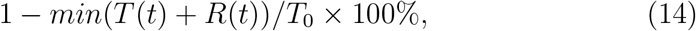

where *T*(*t*) and *R*(*t*) are the number of susceptible and refractory epithelial cells during infection, and *T*_0_ is the initial number of available susceptible cells. The area under the dead cell curve (*AUC_D_*) is given by

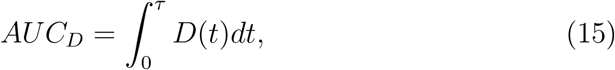

*τ* is a cut-off day for calculation, and we set *τ* = 10 to cover viral and macrophage dynamics shown in [18]. *D*(*t*) is simulated time series of dead cells. The relative contribution of macrophages on viral clearance is given by

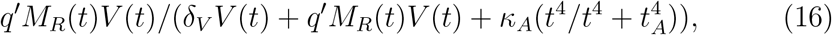

where *M_R_*(*t*) and *V*(*t*) are the number of resting macrophages and viral loads during infection. The prediction of tissue damage and the reproduction number were computed using 6000 posterior samples by solving the ordinary differential equations (ODEs) solver ode15s in MATLAB R2022a with a relative tolerance of 1 × 10^−5^ and an absolute tolerance of 1 × 10^−10^. The initial values were (*M_R_, M*_1_, *M*_2_, *T, L, I, V, F, R, D*) = (*s/δ_MR_*, 0, 0, *T*_0_, 0, 0, *V*_0_, 0, 0, 0). All visualization was performed in R (version 4.0.2), and codes to produce all figures are available at https://github.com/keli5734/virulence.

### A simulation and estimation study

A simulation and estimation study is conducted prior to implement the real dataset. The purpose of the simulation and estimation study is to explore if extra macrophage data provides more information to better estimate model parameters and reproduce viral and macrophage dynamics. We use simulation and mathematical model to show that macrophage data can be used to accurately the recruitment rate of macrophages, inferring the timing and strength of the increase of macrophage during influenza viral infection. By contrast, viral load data alone cannot be used to reliably recover the macrophage dynamics. Hence, the combination of viral load and macrophage data in model fitting enhances our ability to replicate macrophage dynamics and allows us to explore detailed macrophage-virus interactions, e.g., the contribution of macrophages (both in timing and strength) on viral clearance. Detailed study design and outcome can be found in S3 Text in *Supporting Information*.

## Supporting Information

### S1 Text Convergence diagnostics for the MCMC chains

Figures A and B show trace plots for the evolution of estimated parameter vector over the iterations of 3 Markov chains for implementing H1N1 and H5N1 virus, respectively. For each chain, the iteration number is 3000 with the first 1000 samples as burn-in. We observe that all three chains do overlap together, indicating convergence has occurred. Tables A and B show the credible intervals, effective sample size and 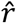 of each estimated parameters for HP and LP strains of H1N1 or H5N1 virus, respectively. We find that the effective sample size is sufficient and 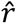 is below 1.1 for every parameter, suggesting convergence.

**Figure A.**
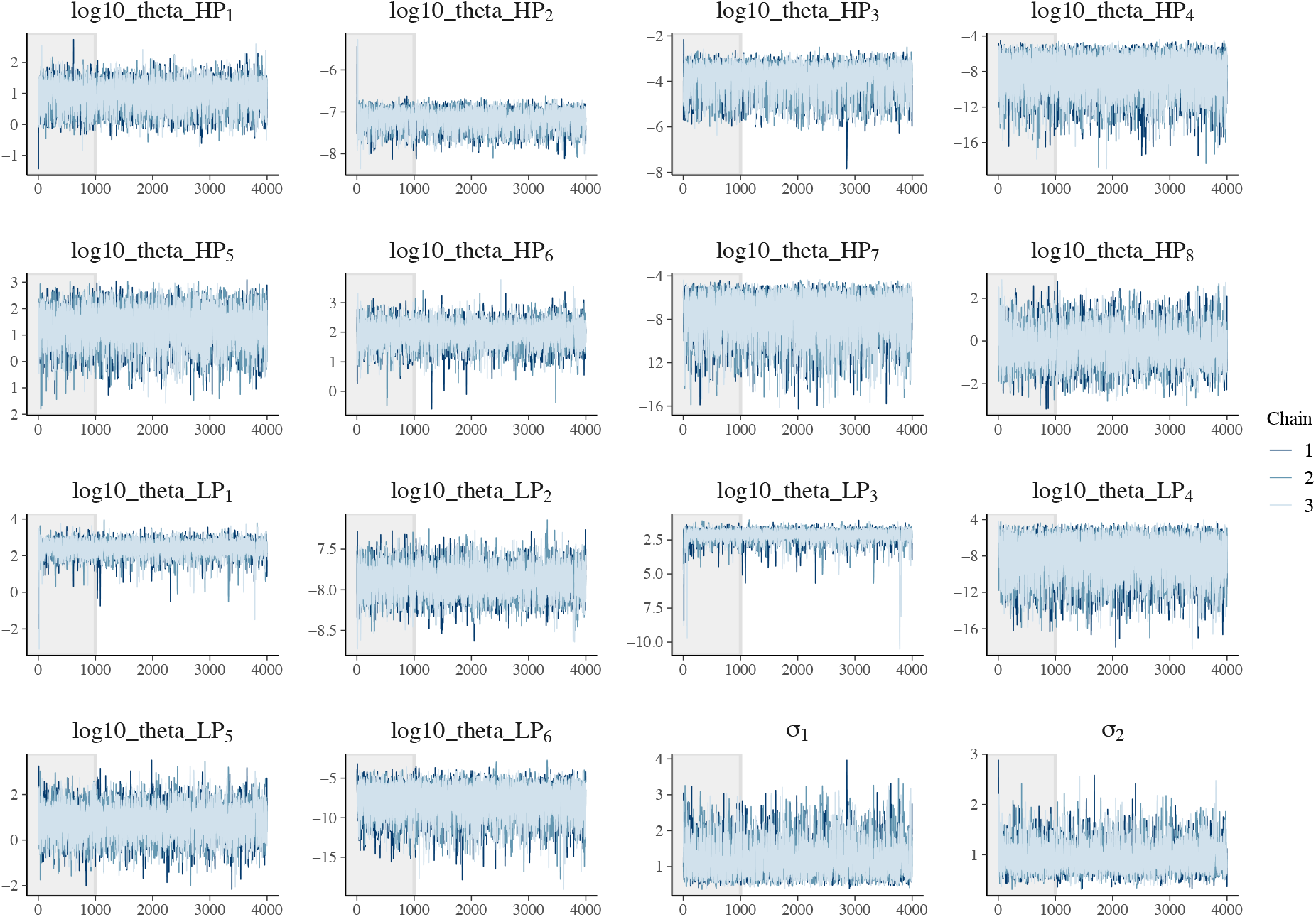
Trace plots of estimated parameters for the fitting H1N1 viral and macrophage data. Three chains were used with 3000 iterations and first 1000 iterations as burn-in (grey area). All parameters are log-transformed. The Parameter vector for HP 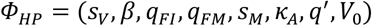, and 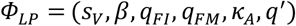 for LP. We assume *s_M_* and *V*_0_ are the same for both HP and LP strains. *σ*_1_, *σ*_2_ are error structure for the prior distribution of standard deviation the observed log-transformed viral load and macrophage data.

**Figure B.**
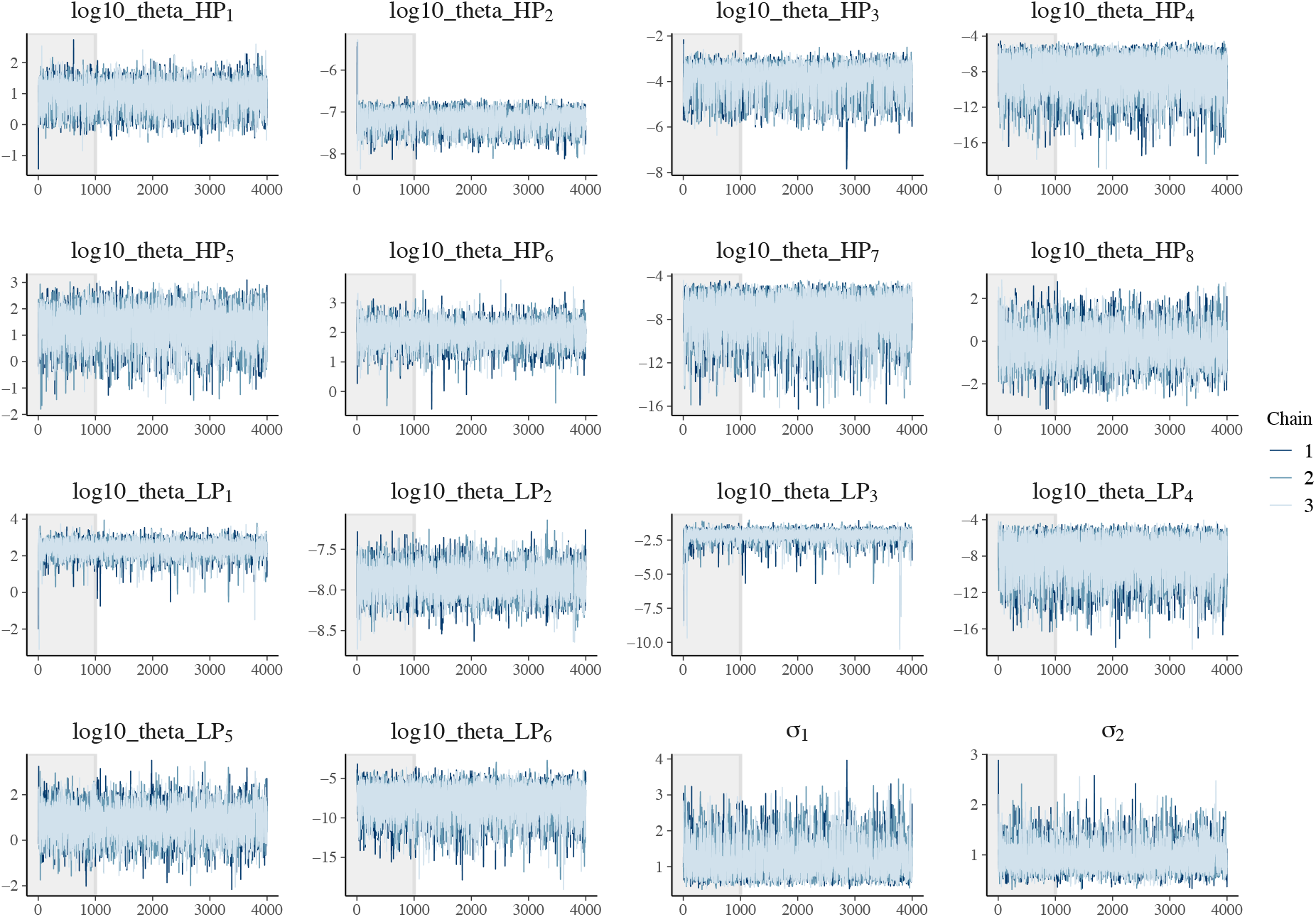
Trace plots of estimated parameters for the fitting H5N1 viral and macrophage data.

**Table A.** Credible intervals, effective sample sizes and 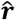 for each estimated parameter of HP and LP strains for H1N1 virus. The first 1000 iterations are discarded as burn-in, leaving 6000 samples across the three chains.

|  | mean | se_mean | sd | 2.5% | 25% | 50% | 75% | 97.5% | n_eff | Rhat |
| --- | --- | --- | --- | --- | --- | --- | --- | --- | --- | --- |
| log10_theta_HP[1] | 0.91 | 0.01 | 0.45 | -0.02 | 0.60 | 0.96 | 1.24 | 1.70 | 1535 | 1.00 |
| log10_theta_HP[2] | -7.20 | 0.01 | 0.23 | -7.70 | -7.35 | -7.15 | -7.03 | -6.84 | 1076 | 1.00 |
| log10_theta_HP[3] | -3.94 | 0.02 | 0.68 | -5.51 | -4.33 | -3.78 | -3.43 | -3.02 | 1014 | 1.01 |
| log10_theta_HP[4] | -8.48 | 0.04 | 2.15 | -13.30 | -9.81 | -8.19 | -6.80 | -5.34 | 3351 | 1.00 |
| log10_theta_HP[5] | 1.33 | 0.01 | 0.69 | -0.21 | 0.92 | 1.38 | 1.81 | 2.51 | 2724 | 1.00 |
| log10_theta_HP[6] | 1.97 | 0.01 | 0.37 | 1.14 | 1.76 | 1.99 | 2.20 | 2.66 | 3152 | 1.00 |
| log10_theta_HP[7] | -7.82 | 0.04 | 2.00 | -12.43 | -9.07 | -7.51 | -6.21 | -5.02 | 2428 | 1.00 |
| log10_theta_HP[8] | -0.17 | 0.02 | 0.84 | -1.78 | -0.75 | -0.21 | 0.38 | 1.53 | 1486 | 1.00 |
| log10_theta_LP[1] | 2.36 | 0.01 | 0.43 | 1.39 | 2.16 | 2.40 | 2.62 | 3.06 | 1499 | 1.00 |
| log10_theta_LP[2] | -7.89 | 0.00 | 0.18 | -8.26 | -8.01 | -7.89 | -7.77 | -7.55 | 2026 | 1.00 |
| log10_theta_LP[3] | -2.21 | 0.03 | 0.56 | -3.31 | -2.37 | -2.11 | -1.92 | -1.60 | 438 | 1.00 |
| log10_theta_LP[4] | -8.27 | 0.04 | 2.23 | -13.26 | -9.68 | -7.98 | -6.50 | -4.98 | 3159 | 1.00 |
| log10_theta_LP[5] | 0.86 | 0.02 | 0.69 | -0.58 | 0.44 | 0.88 | 1.31 | 2.18 | 2055 | 1.00 |
| log10_theta_LP[6] | -7.92 | 0.04 | 2.03 | -12.67 | -9.08 | -7.60 | -6.44 | -4.85 | 2883 | 1.00 |
| sigma[1] | 1.24 | 0.01 | 0.48 | 0.58 | 0.88 | 1.13 | 1.51 | 2.39 | 1047 | 1.00 |
| sigma[2] | 0.93 | 0.01 | 0.27 | 0.52 | 0.74 | 0.89 | 1.08 | 1.58 | 1815 | 1.00 |

**Table B.** Credible intervals, effective sample sizes and 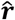 for each estimated parameter of HP and LP strains for H5N1 virus.

|  | mean | se_mean | sd | 2.5% | 25% | 50% | 75% | 97.5% | n_eff | Rhat |
| --- | --- | --- | --- | --- | --- | --- | --- | --- | --- | --- |
| log10_theta_HP[1] | 0.87 | 0.01 | 0.51 | -0.08 | 0.52 | 0.89 | 1.22 | 1.81 | 1613 | 1.00 |
| log10_theta_HP[2] | -7.24 | 0.01 | 0.24 | -7.72 | -7.41 | -7.23 | -7.07 | -6.82 | 1859 | 1.00 |
| log10_theta_HP[3] | -4.01 | 0.02 | 0.69 | -5.52 | -4.45 | -3.88 | -3.49 | -2.97 | 1617 | 1.00 |
| log10_theta_HP[4] | -8.45 | 0.04 | 2.17 | -13.37 | -9.81 | -8.09 | -6.78 | -5.35 | 3277 | 1.00 |
| log10_theta_HP[5] | 1.16 | 0.01 | 0.70 | -0.32 | 0.71 | 1.21 | 1.66 | 2.39 | 3527 | 1.00 |
| log10_theta_HP[6] | 1.99 | 0.02 | 0.53 | 0.91 | 1.72 | 1.98 | 2.24 | 3.18 | 649 | 1.00 |
| log10_theta_HP[7] | -7.87 | 0.04 | 1.99 | -12.57 | -9.08 | -7.54 | -6.36 | -5.03 | 2456 | 1.00 |
| log10_theta_HP[8] | 0.22 | 0.02 | 0.91 | -1.59 | -0.39 | 0.22 | 0.82 | 2.03 | 2317 | 1.00 |
| log10_theta_LP[1] | 2.16 | 0.03 | 0.64 | 0.66 | 1.87 | 2.24 | 2.54 | 3.11 | 616 | 1.00 |
| log10_theta_LP[2] | -7.87 | 0.01 | 0.24 | -8.36 | -8.03 | -7.85 | -7.69 | -7.44 | 730 | 1.00 |
| log10_theta_LP[3] | -2.98 | 0.06 | 1.12 | -6.47 | -3.12 | -2.73 | -2.43 | -1.93 | 330 | 1.01 |
| log10_theta_LP[4] | -8.41 | 0.04 | 2.10 | -13.03 | -9.78 | -8.11 | -6.77 | -5.35 | 3412 | 1.00 |
| log10_theta_LP[5] | 1.06 | 0.01 | 0.71 | -0.46 | 0.62 | 1.10 | 1.54 | 2.37 | 2256 | 1.00 |
| log10_theta_LP[6] | -8.08 | 0.04 | 2.00 | -12.61 | -9.23 | -7.76 | -6.61 | -5.04 | 2715 | 1.00 |
| sigma[1] | 1.74 | 0.02 | 0.50 | 0.81 | 1.40 | 1.72 | 2.05 | 2.80 | 918 | 1.00 |
| sigma[2] | 0.93 | 0.02 | 0.38 | 0.47 | 0.67 | 0.82 | 1.07 | 1.96 | 416 | 1.01 |

### S2 Text Parameter tables

#### Fixed Parameter table (Table S1)

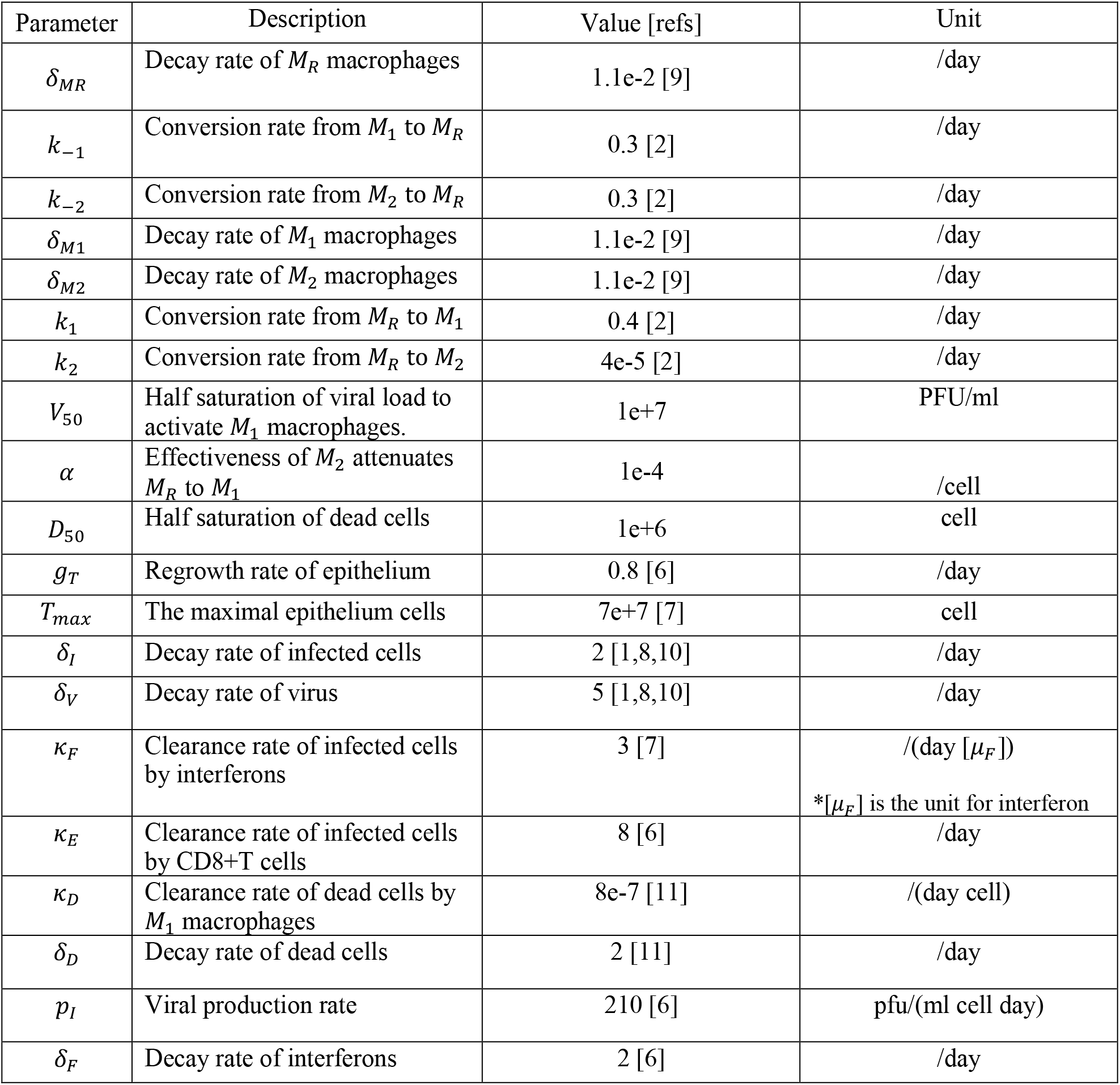
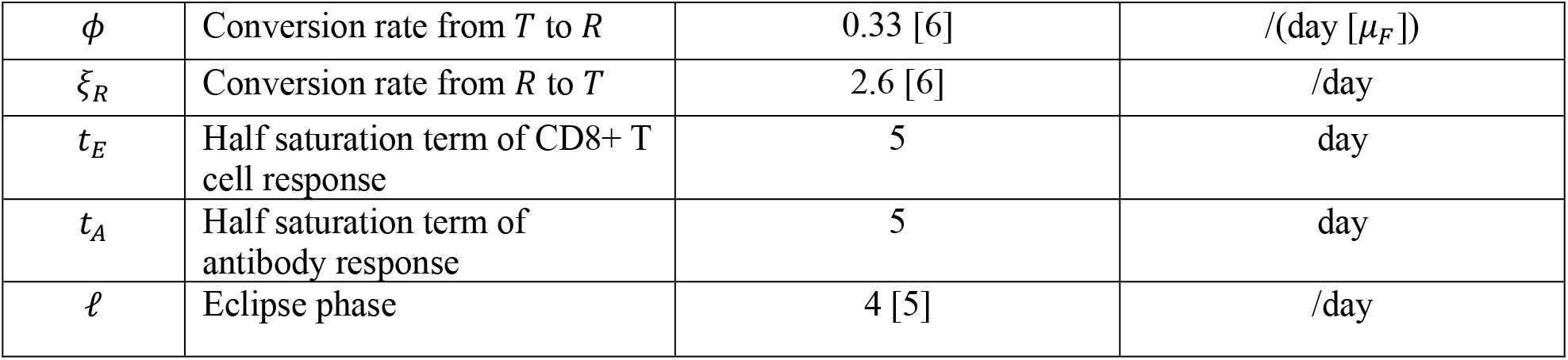

#### Estimated Parameter table (Table S2)

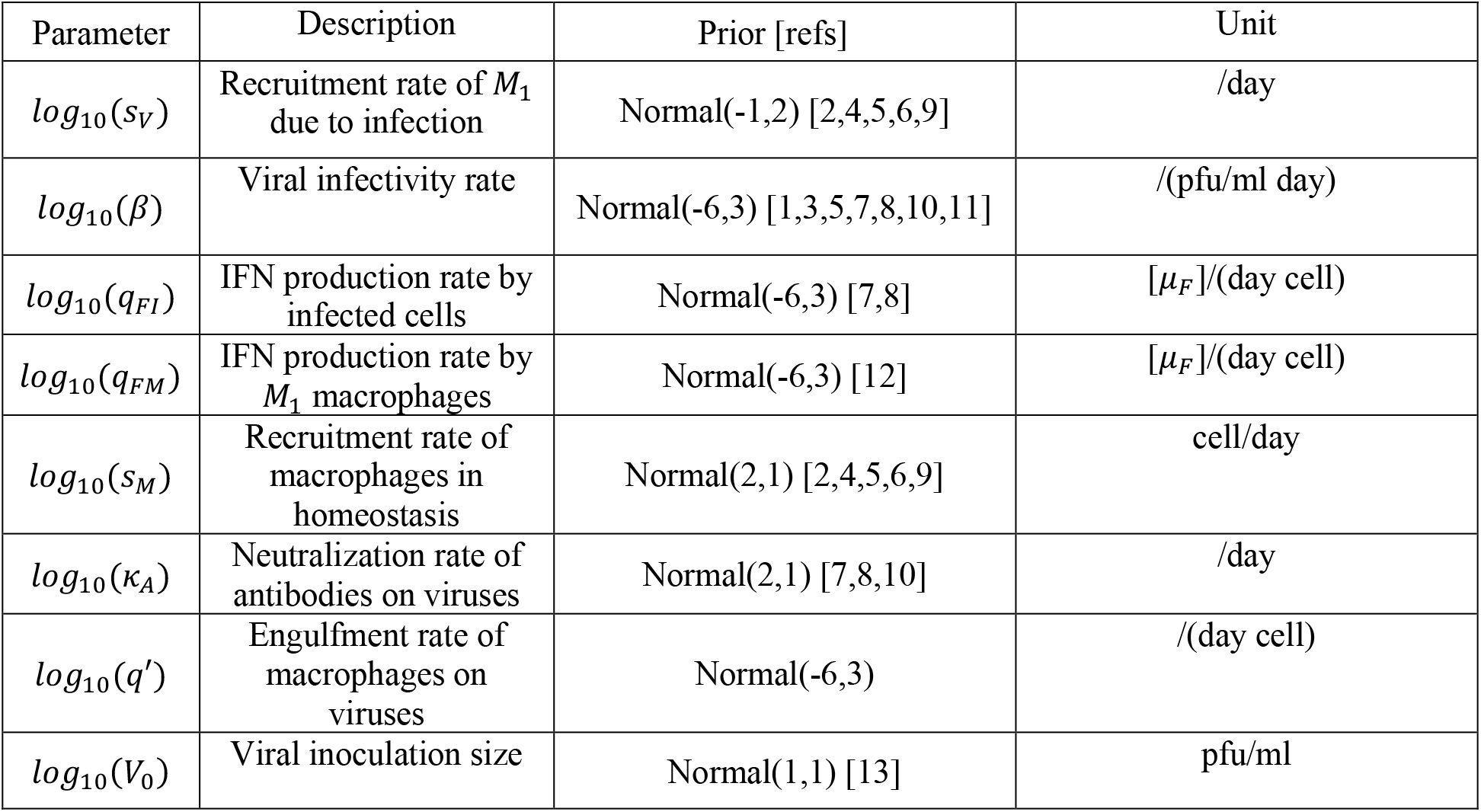

## S3 Text Simulation and estimation study

The purpose of the simulation and estimation study is to explore if extra macrophage data provides more information to better estimate model parameters and reproduce viral and macrophage dynamics. We use simulation and mathematical model to show that macrophage data can be used to accurately the recruitment rate of macrophages, inferring the timing and strength of the increase of macrophage during influenza viral infection. By contrast, viral load data alone cannot be used to reliably recover the macrophage dynamics. Hence, the combination of viral load and macrophage data in model fitting enhances our ability to replicate macrophage dynamics and allows us to explore detailed macrophage-virus interactions, e.g., the contribution of macrophages (both in timing and strength) on viral clearance.

### Generation of synthetic viral load and macrophage data

We first generate synthetic data for viral loads and macrophages mimicking the experimental procedure. We assume “true” parameter values are known (see Table 1). We do model (details in main text) simulation using the parameter values to get “true” trajectory of viral load and macrophages dynamics across infection period. The “true” parameters are selected such that (1) viral load peaks around day 2 post infection; (2) viral load is below a detection limit around day 7 post infection; (3) the adaptive immune responses (i.e., antibody and CD8+T cells) only activate after day 5 post infection; (4) viral infection can be suppressed timely when both arms of adaptive immune responses (i.e., antibody and CD8+T cells) are presented; (5) virus can be cleared but clearance delays when there is only an antibody response, and (6) a chronic infection occurs when an antibody repones is suppressed (Figure 1C). A detailed model dynamics see Figure 1.

Further, we get observation viral load and macrophage data from the “true” trajectory by adding lognormal noise and imposing a detection limit. Mathematically, the measured viral load *V_n,τ_* and macrophage *M_n,τ_* for each mouse *n* = 1,2, …, *N* and measuring time point *τ* = 1, 2, …, *T* are given by

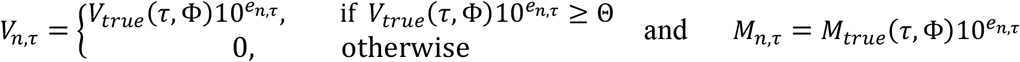

Φ is a vector of “true” parameter values. *e_n,τ_* is the measurement error, which follows *N*(0, *σ*), and *σ* = 1 for viral load data and *σ* = 0.1 for macrophage data. Θ is detection limit. *V_true_*(*τ*, Φ) is the “true” viral load value at each measuring time *τ*, and *M_true_*(*τ*, Φ) is the “true” macrophage value at each measuring time *τ*. Here, we select *N* = 5 to indicate at each measuring time 5 data points are measured, and we set *τ* = 7. As shown in Figures 1A and B, the open circles indicate measured data points at each measuring time for viral load and macrophages, respectively. The red cycles indicate the mean value of the 5 data point at each time, and we only use the red data points of viral load and macrophage populations for the model estimation.

**Figure 1.**
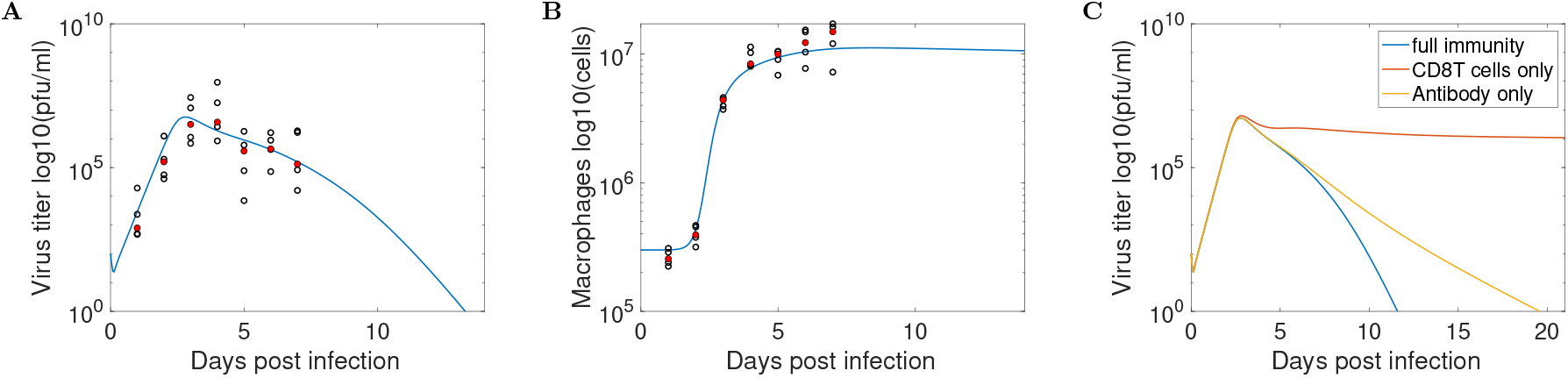
The synthetic data for (A) viral load, (B) macrophages. (C) “true” parameter values are selected such that viral loads have different behaviours when different arms of adaptive immune responses are suppressed.

**Table 1:** “true” parameter values to generate true viral load and macrophages trajectories

| Parameter | Description | Value | Unit |
| --- | --- | --- | --- |
| $\delta_{MR}$ | Decay rate of $M_R$ macrophages | 3.3e+3 | /day |
| $k_{-1}$ | Conversion rate from $M_1$ to $M_R$ | 0.3 | /day |
| $k_{-2}$ | Conversion rate from $M_2$ to $M_R$ | 0.3 | /day |
| $\delta_{M1}$ | Decay rate of $M_1$ macrophages | 1.1e-2 | /day |
| $\delta_{M2}$ | Decay rate of $M_2$ macrophages | 1.1e-2 | /day |
| $k_1$ | Conversion rate from $M_R$ to $M_1$ | 0.4 | /day |
| $k_2$ | Conversion rate from $M_R$ to $M_2$ | 4e-6 | /day |
| $V_{50}$ | Half saturation of viral load to activate $M_1$ macrophages. | 1e+7 | PFU/ml |
| $\alpha$ | Effectiveness of $M_2$ attenuates $M_R$ to $M_1$ | 1e-4 | /cell |
| $D_{50}$ | Half saturation of dead cells | 1e+6 | cell |
| $g_T$ | Regrowth rate of epithelium | 0.8 | /day |
| $T_{max}$ | The maximal epithelium cells | 7e+7 | cell |
| $\delta_I$ | Decay rate of infected cells | 2 | /day |
| $\delta_V$ | Decay rate of virus | 20 | /day |
| $\kappa_F$ | Clearance rate of infected cells by interferons | 3 | /(day [$\mu_F$])<br>*$[\mu_F]$ is the unit for interferon |
| $\kappa_E$ | Clearance rate of infected cells by CD8+T cells | 8 | /day |
| <sup>455</sup>$\kappa_D$ | Clearance rate of dead cells by $M_1$ macrophages | 8e-7 | /(day cell) |
| $\delta_D$ | Decay rate of dead cells | 2 | /day |
| $p_I$ | Viral production rate | 210 | pfu/(ml cell day) |
| $\delta_F$ | Decay rate of interferons | 2 | /day |
| $\phi$ | Conversion rate from $T$ to $R$ | 0.33 | /(day [$\mu_F$]) |
| $\xi_R$ | Conversion rate from $R$ to $T$ | 2.6 | /day |
| $\ell$ | Eclipse phase | 4 | /day |
| $t_E$ | Half saturation time of CD8+ T cell response | 5 | day |
| $t_A$ | Half saturation time of antibody response | 5 | day |

### Bayesian statistical inference

Two scenarios were considered, one of which is using viral load data only, and the other is using both viral load and macrophage data. We applied a Bayesian inference method to fit the dynamic model (detailed in the main text) to the log-transformed kinetic data. In detail, we use the model to estimate 8 parameters, and the parameter space is denoted as 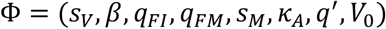. Upon model calibration, we fixed all other parameters their “true” values as shown in Table 1.

The prior distribution for the estimated model parameters is given in Table S2 in Supplementary Materials 2. The distribution of the observed log-transformed viral load and/or macrophage data is assumed to be a normal distribution with a mean value given by the model simulation results and standard deviation (SD) parameter with prior distribution of a normal distribution with a mean of 0 and a SD of 1.

Model fitting was performed in R (version 4.0.2) and Stan (Rstan 2.21.0). Samples were drawn from the joint posterior distribution of the model parameters using Hamiltonian Monte Carlo (HMC) optimized by the No-U-Turn Sampler (NUTS) (details see Chatzilena et al. (2019)). In particular, we used 4 chains with different starting points and ran 8000 iterations (first 3000 samples are burn-in) for each chain when only viral load data is used. We also tried to run 2000, 4000 and 6000 iterations, respectively and effective sample size is small. When viral load data and macrophages are both used, we ran 4 chains with 2000 iterations (first 1000 samples are burn-in) for each chain.

### Predictive check

**Figure 2.**
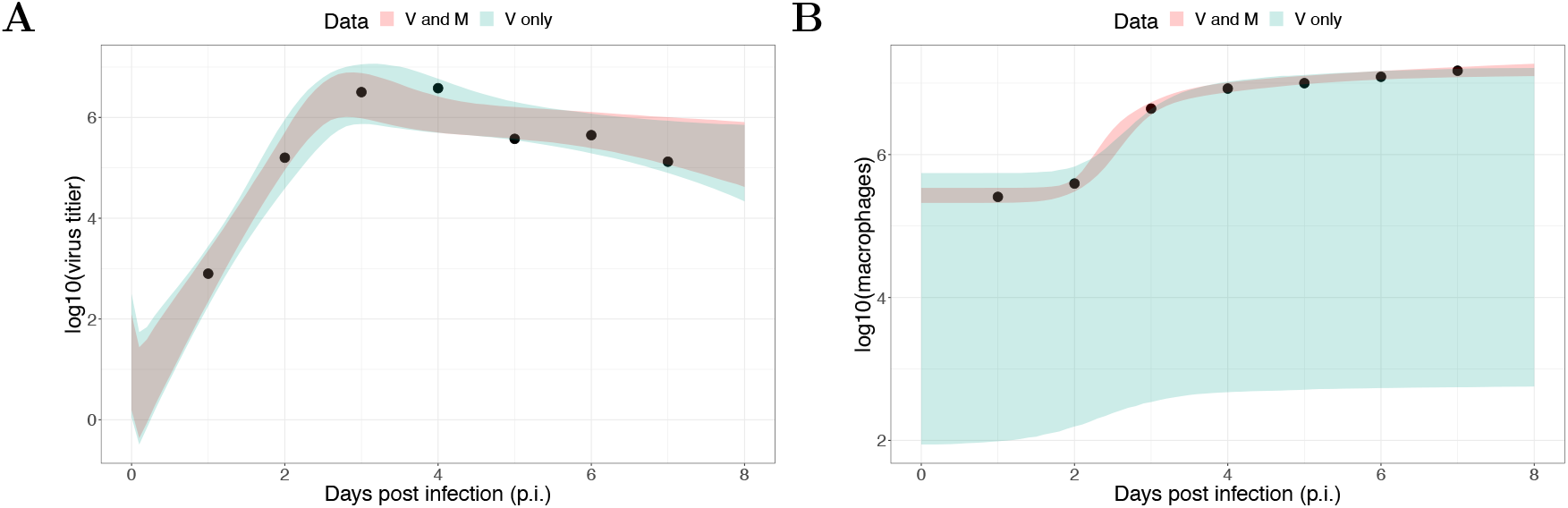
Results of model fitting for virological and macrophage data. Data are presented by solid circles. (A) shows a 95% prediction interval (shaded area) of reproduced viral dynamics by using viral load data only (red) or both viral load and macrophage data (green). (B) shows a 95% prediction interval (shaded area) of reproduced macrophage kinetics by using viral load data only (red) or both viral load and macrophage data (green).

### Posterior comparison

**Figure 3.**
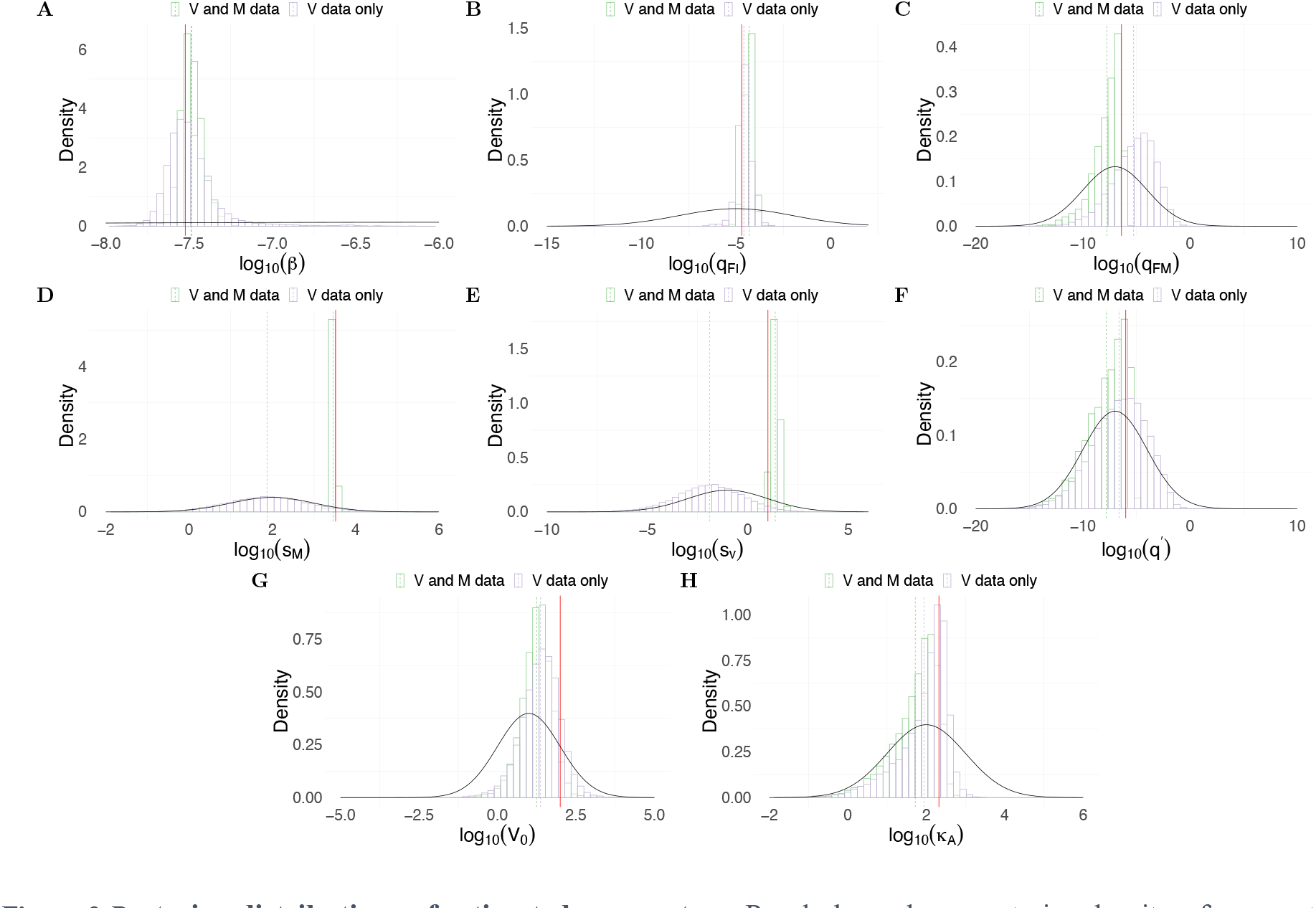
Posterior distributions of estimated parameters. Purple bars show posterior density of parameters when only viral load data is used. Green bars show posterior density of parameters when both viral load and macrophage data are used. Red lines indicate the “true” parameter values.

### Diagnostics

**Figure 4.**
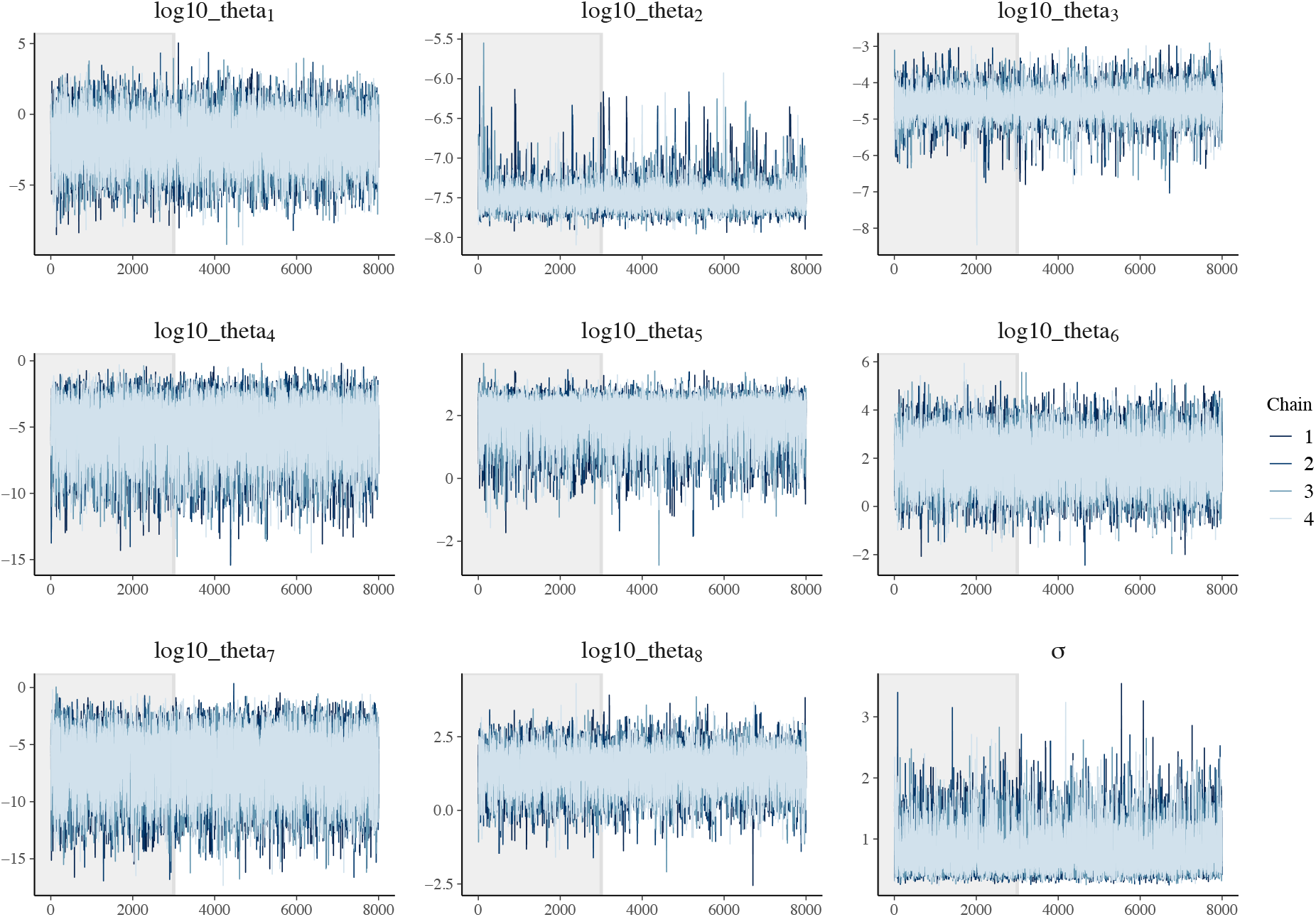
Trace plots of estimated parameters using only viral load data. Four chains were used with 8000 iterations and first 3000 iterations as burn-in (grey area). All parameters are log-transformed. The parameter vector is 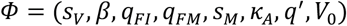.

**Figure 5.**
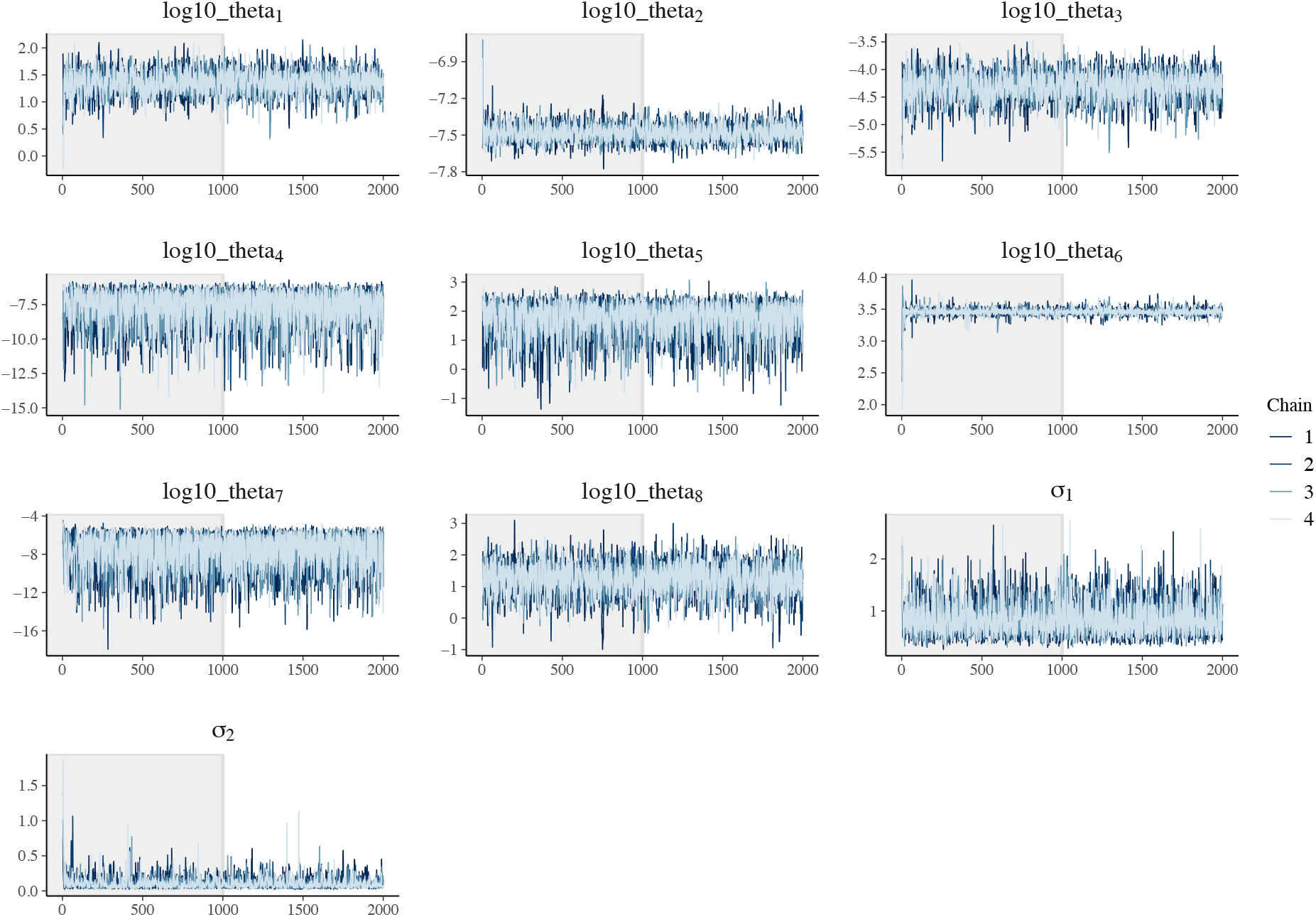
Trace plots of estimated parameters using both viral load and macrophage data. Four chains were used with 2000 iterations and first 1000 iterations as burn-in (grey area). All parameters are log-transformed. The parameter vector is 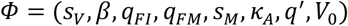.

## Supplementary Figures

**SFig 1.**
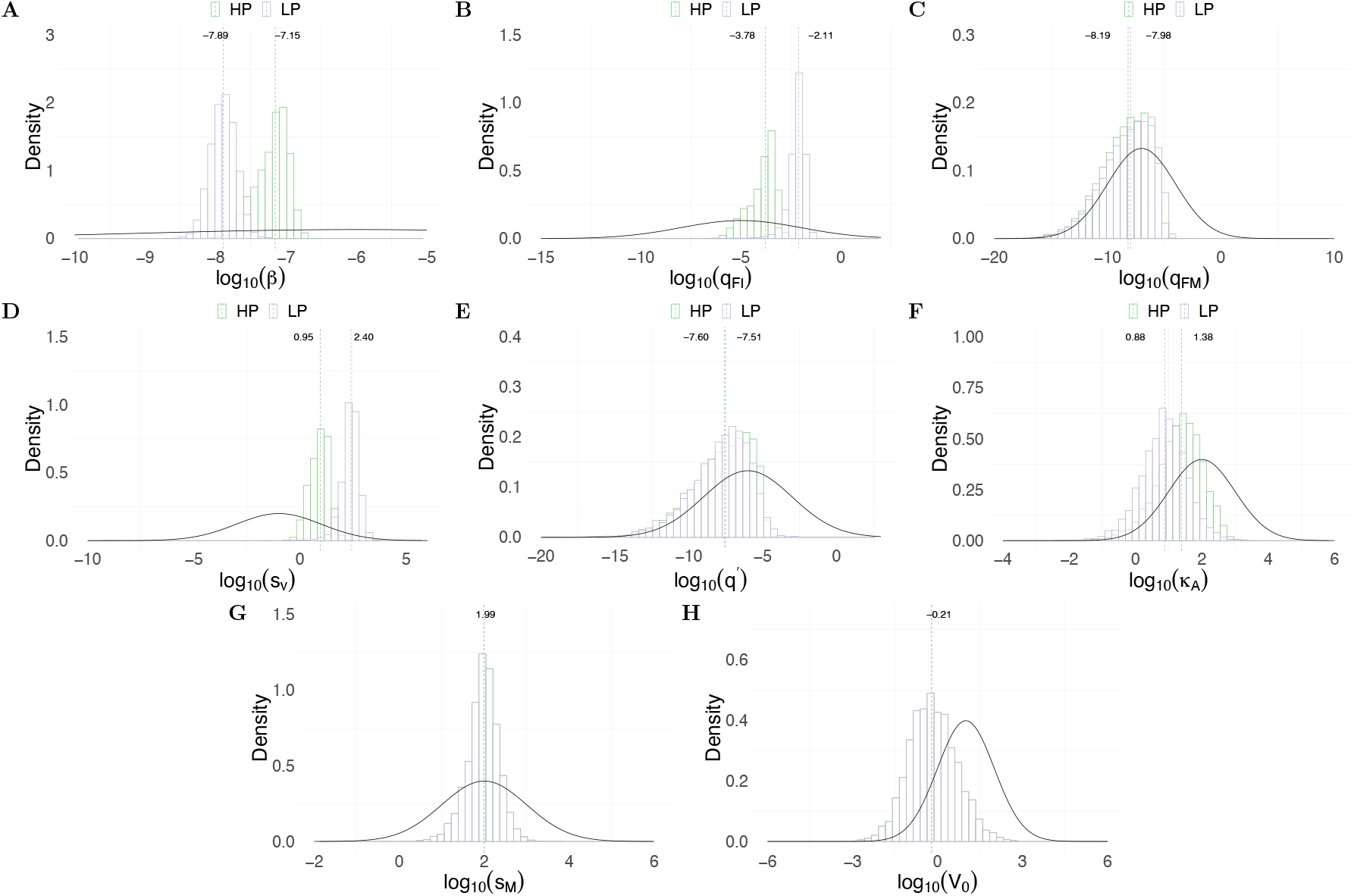
Posterior distributions of parameters for H1N1 virus. Green bars indicate the posterior density for HP strain and purple bars indicate the posterior density for LP strain. Green and purple dashed lines indicate the median estimation of each parameter for HP and LP, respectively. Prior distribution for each parameter is given by black curve.

**SFig 2.**
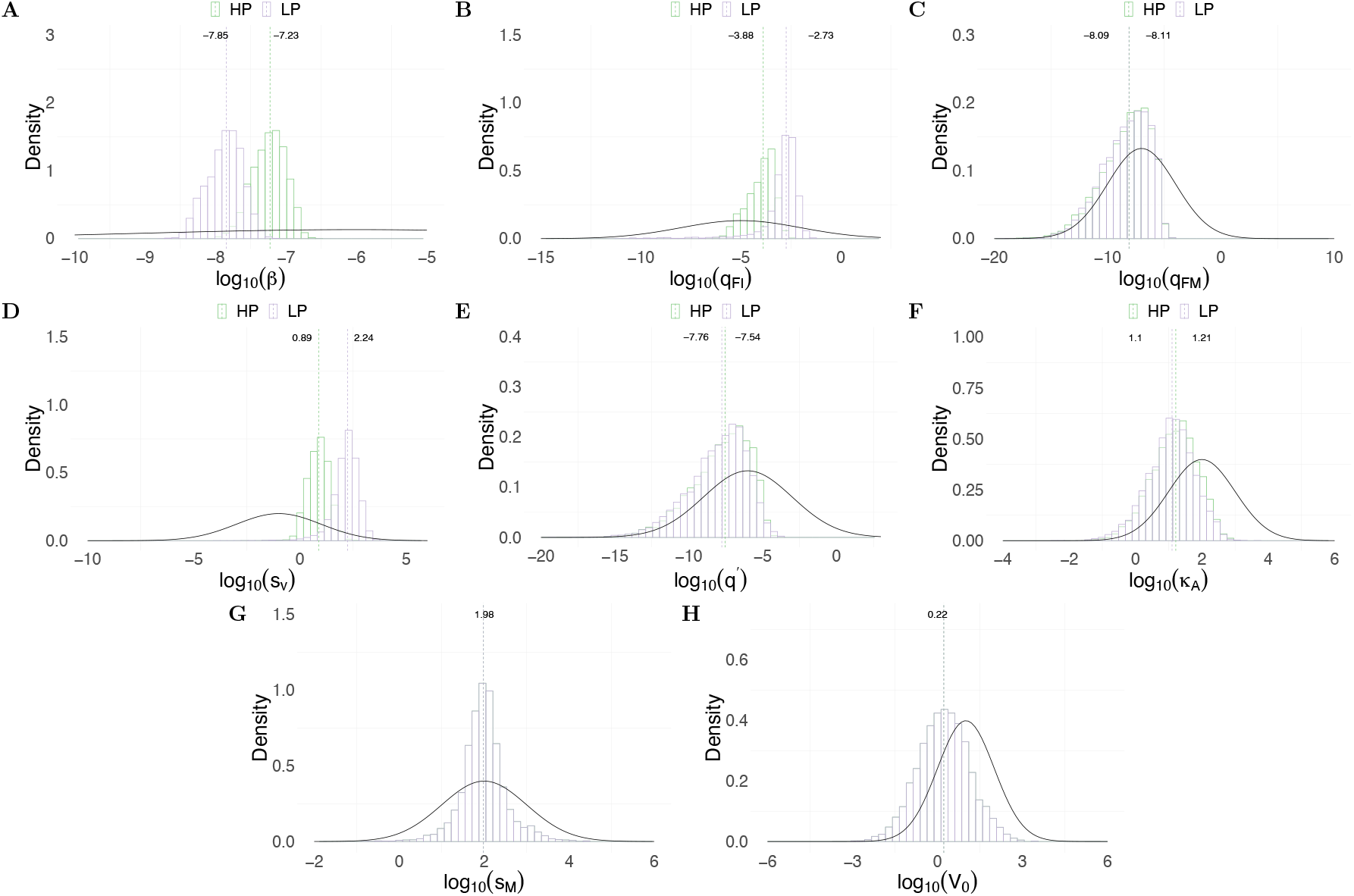
Posterior distributions of parameters for H5N1 virus. Green bars indicate the posterior density for HP strain and purple bars indicate the posterior density for LP strain. Green and purple dashed lines indicate the median estimation of each parameter for HP and LP, respectively. Prior distribution for each parameter is given by black curve.

**SFig 3.**
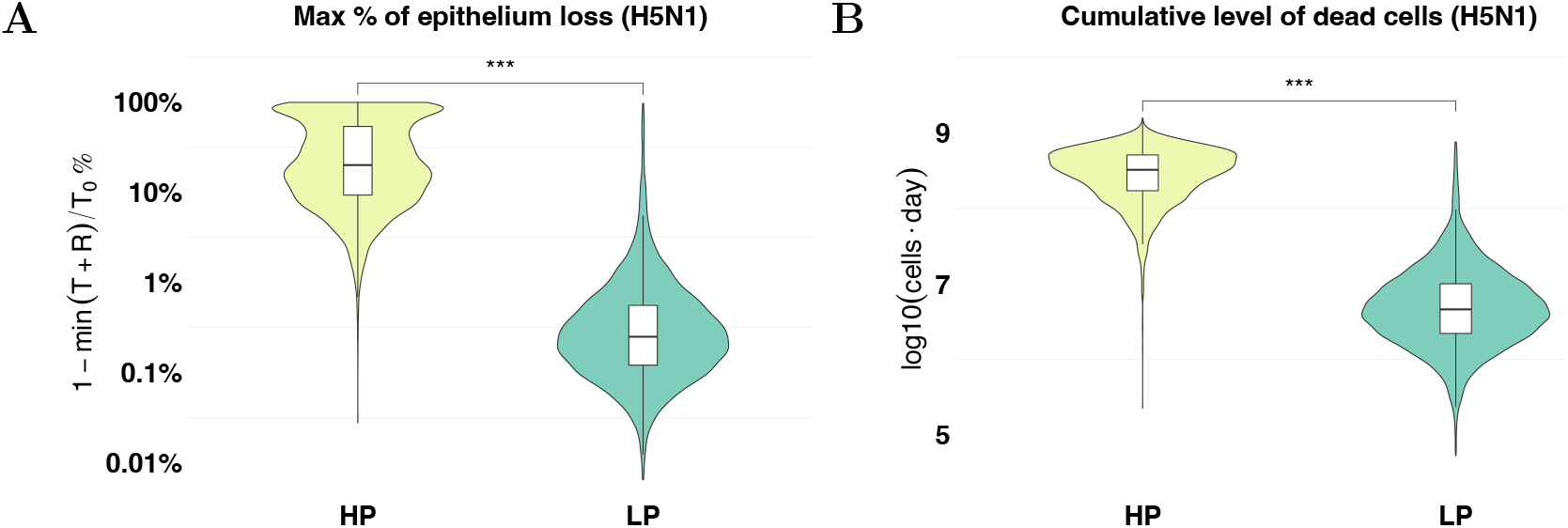
Prediction of tissue damage for H5N1 viruses. The violin plots (coloured) and boxplots (white) give the density and the median and extrema of predicted quantity. (A) model prediction of the maximal epithelium loss for the HP (yellow) and green (LP) strain. (B) model prediction of the cumulative level of dead cells during the infection for both strains. *** *p* < 0.001. Calculation formula see Eq. (13) in the main text. All estimations are computed using 6000 posterior samples from model fitting.

**SFig 4.**
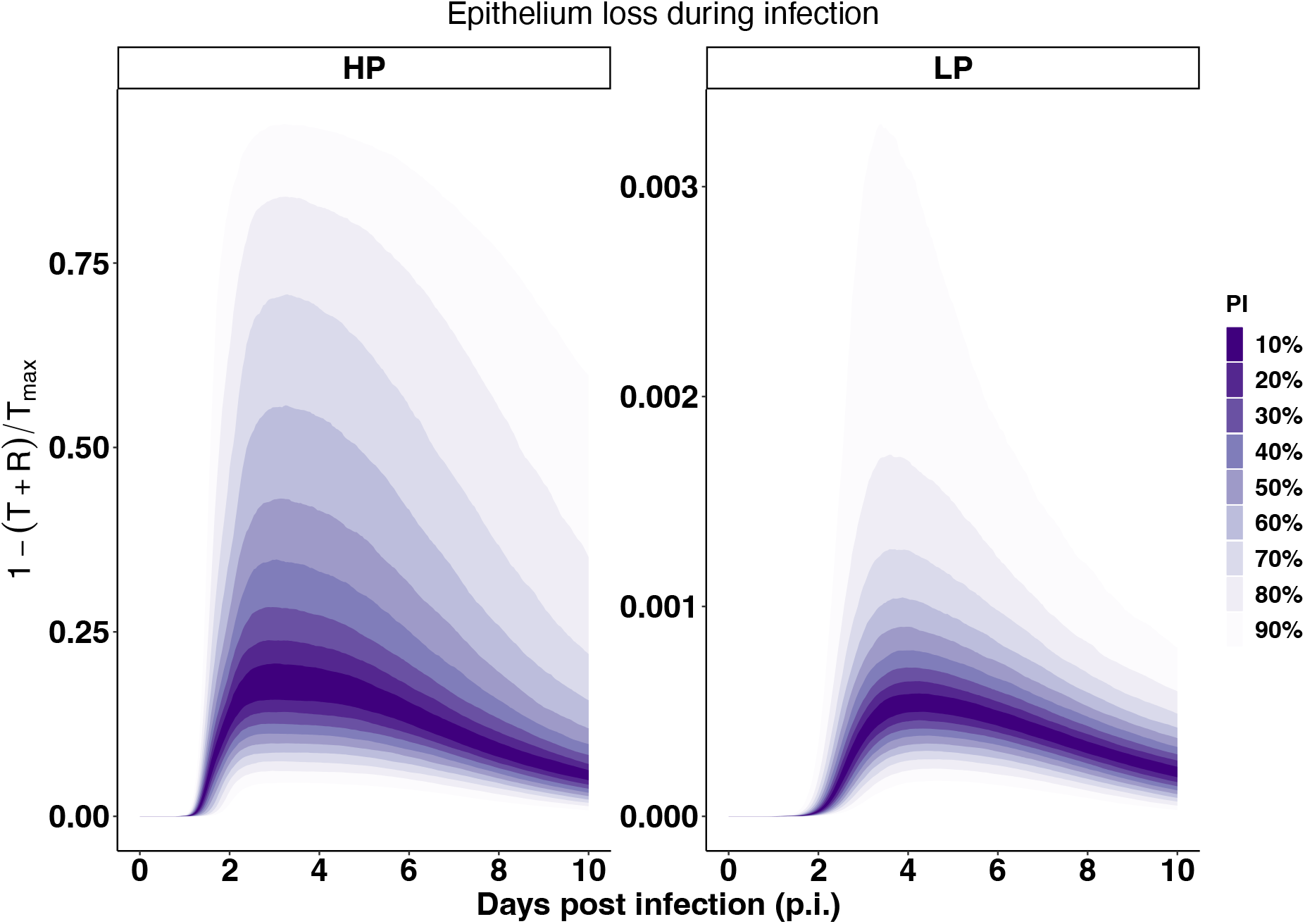
The proportional of epithelium loss during HP and LP H1N1 viral infections. The calculation of epithelium loss is given in the main text. All estimations are computed using 6000 posterior samples from model fitting.

**SFig 5.**
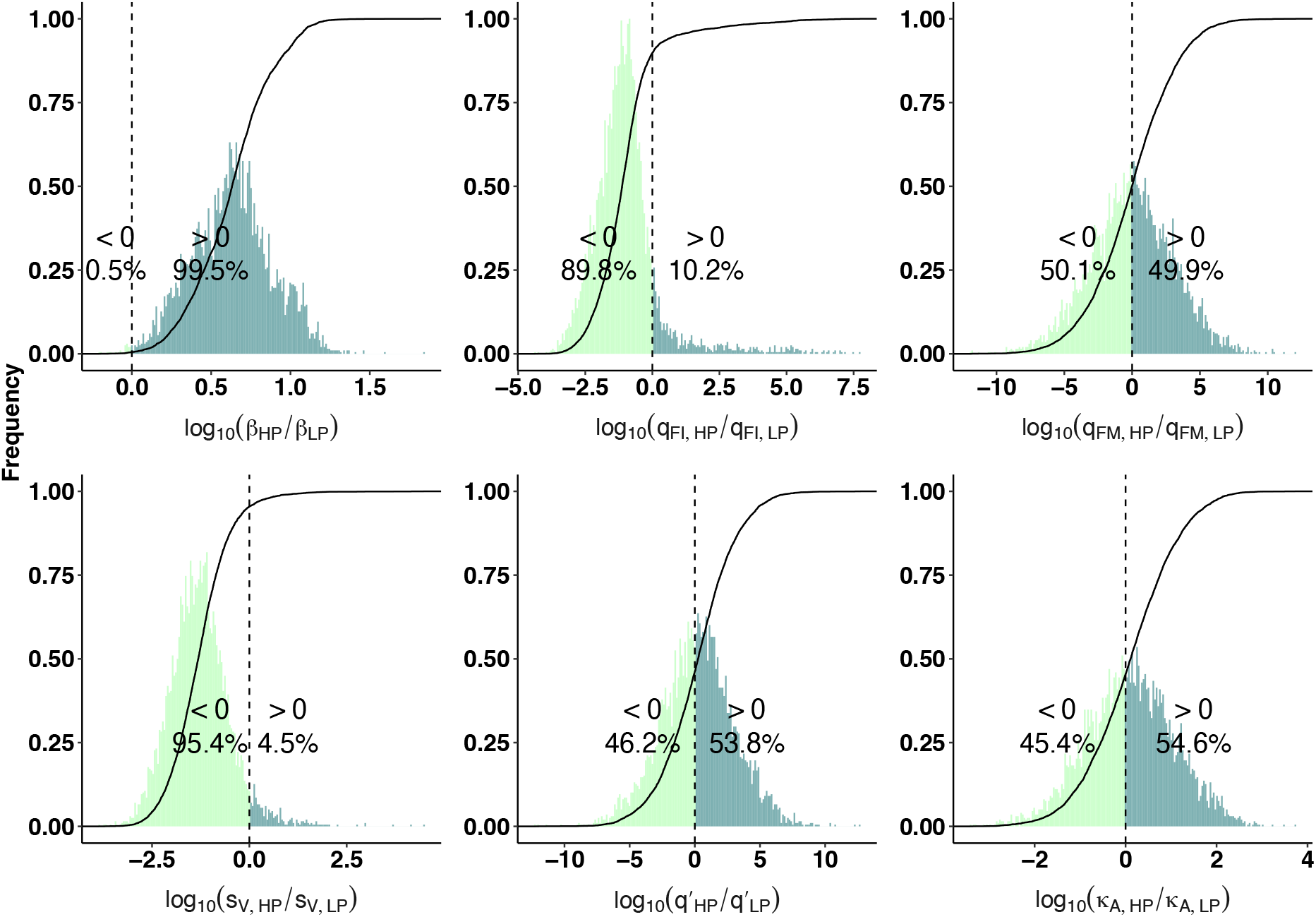
Comparison of estimated model parameters between HP and LP strains of the H5N1 viruses. Histograms show the frequency of the quotient of estimated HP parameters over paired LP model parameters and are normalised to [0,1]. The cumulative density functions (CDF) are given by the solid lines. All quotients are log10-scaled, such that quotient > 0 suggests greater values of the HP parameters. Dark green indicates quotients > 0, and light green indicates quotients < 0. First row (from left to right) the quotients of viral infectivity, interferon production rate from infected cells and activated macrophages, respectively. Second row (from left to right) the quotients of infection-induced macrophage recruitment rate, macrophage-mediated virus clearance rate and antibody neutralisation rate, respectively.

**SFig 6.**
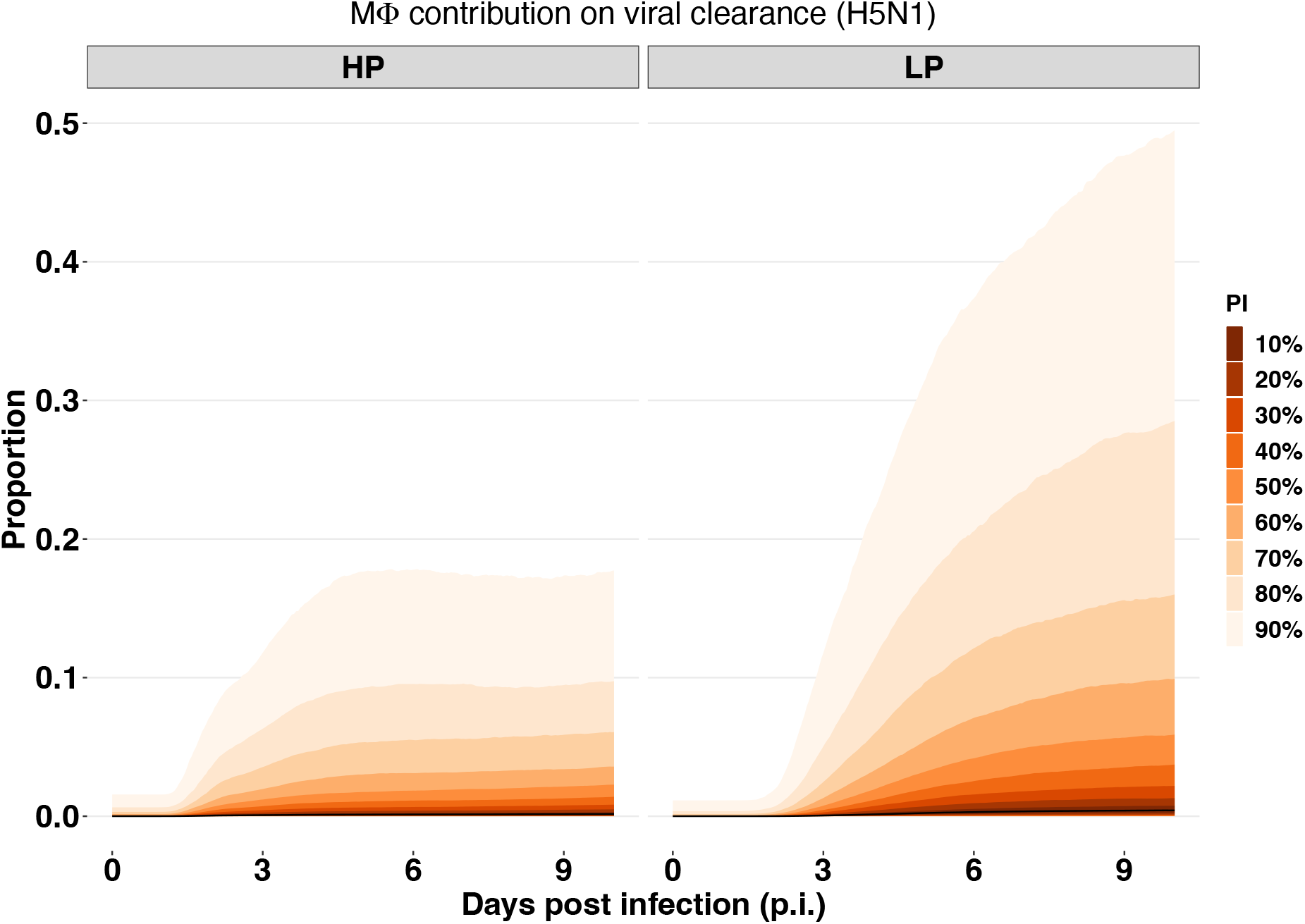
The relative contribution of macrophages on viral clearance in the HP and LP strains of the H5N1 viruses. The relative contribution is given by 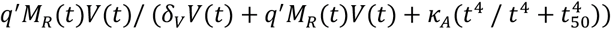, where *M_R_*(*t*) and *V*(*t*) are the number of resting macrophages and viral loads during infection. The prediction interval (PI) is calculated based upon the 6000 posterior samples from model fitting. The median trajectory is indicated by black curve.

**SFig 7.**
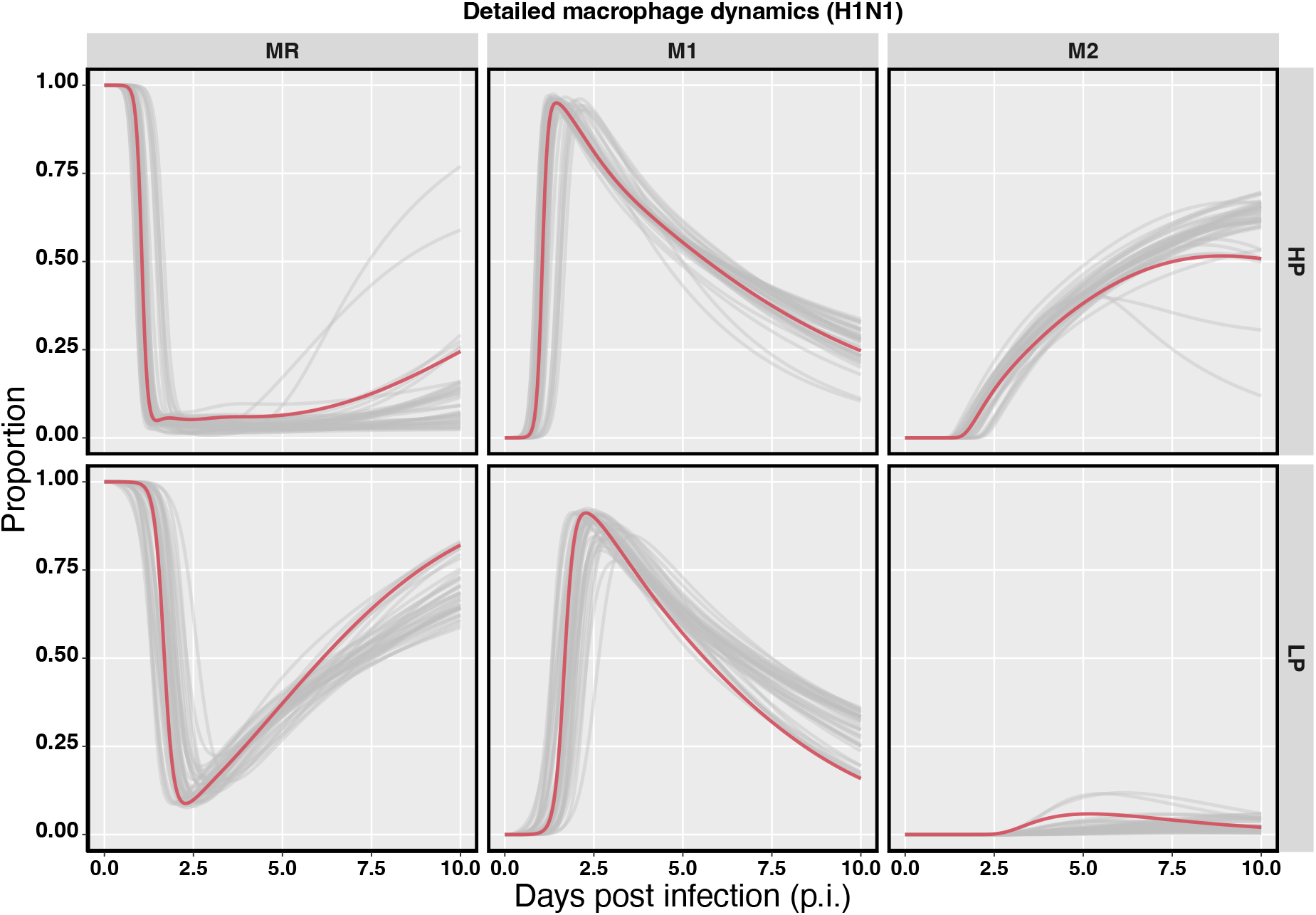
Detailed macrophage dynamics during HP and LP H1N1 viral infections. Y-axis gives the proportion of each type of macrophages to overall number of macrophages at each measuring time. Grey lines are macrophage trajectories calculated based upon 6000 posterior samples from model fitting, and the median trajectory is indicated by red curve.

## Acknowledgements

Ke Li is supported by a Melbourne Research Scholarship. This work was supported by an Australian Research Council (ARC) Discovery Project (DP170103076 and DP210101920) and a National Health and Medical Research Council (NHMRC) funded Centre for Research Excellence in Infectious Diseases Modelling to Inform Public Health Policy (1078068).

## Author contributions

**Ke Li:** Conceptualization, Methodology, Software, Formal analysis, Writing-Original Draft. **James M. McCaw** Methodology, Formal analysis, Writing-Review and Editing, Supervision. **Pengxing Cao:** Methodology, Formal analysis, Writing-Review and Editing, Supervision

